# Genome-wide identification and characterization of major RNAi genes highlighting their regulatory factors in Cowpea (*Vigna unguiculata* (L.) Walp.)

**DOI:** 10.1101/2023.02.15.528631

**Authors:** Mohammad Nazmol Hasan, Md Parvez Mosharaf, Khandoker Saif Uddin, Keya Rani Das, Nasrin Sultana, Mst. Noorunnahar, Darun Naim, Md. Nurul Haque Mollah

## Abstract

In different regions of the world, cowpea (Vigna unguiculata (L.) Walp.) is an important vegetable and an excellent source of protein. It lessens the malnutrition of the underprivileged in developing nations and has some positive effects on health, such as a reduction in the prevalence of cancer and cardiovascular disease. However, occasionally, certain biotic and abiotic stresses caused a sharp fall in cowpea yield. Currently, short RNAs (sRNAs) are thought to play a crucial role in controlling how stress-related genes are expressed in plants as they go through various developmental phases. Major RNA interference (RNAi) genes like Dicer-like (DCL), Argonaute (AGO), and RNA-dependent RNA polymerase (RDR) are essential for sRNA synthesis processes and shield plants from biotic and abiotic stresses. In this study, applying BLASTP search and phylogenetic tree analysis with reference to the *Arabidopsis* RNAi (AtRNAi) genes, we discovered 28 VuRNAi genes, including 7 *VuDCL*, 14 *VuAGO*, and 7 *VuRDR* genes in cowpea. We looked at the domains, motifs, gene structures, chromosomal locations, sub-cellular locations, gene ontology (GO) terms and regulatory factors (transcription factors, micro-RNAs, and *cis*-acting factors) to characterize the VuRNAi genes and proteins in cowpea. Predicted *VuDCL1*, *VuDCL2*(a, b), *VuAGO7*, *VuAGO10* and *VuRDR6* genes might have an impact on cowpea growth, development of the vegetative and flowering stages and antiviral defense. The VuRNAi gene regulatory features miR395 and miR396 might contribute to grain quality improvement, immunity boosting, and pathogen infection resistance under salinity and drought conditions. Predicted *cis*-acting elements (CAEs) from the VuRNAi gene associated with light responsiveness (LR), stress responsiveness (SR) and hormone responsiveness (HR) might play a role in plant growth and development, improving grain quality and production and protecting plants from biotic and abiotic stresses. Therefore, our study provides crucial information about the functional roles of VuRNAi genes and their regulatory components, which would aid in the development of future cowpeas that are more resilient to biotic and abiotic stress.

## 1 Introduction

The cowpea (*Vigna unguiculata* (L.) Walp.) is a significant food and protein source for thousands of people in diverse parts of the world, including tropical Africa, Asia and North and South America, where it is mostly grown. It is also used as a vegetable and is among China’s top ten most significant veggies. Additionally, it contributes significantly to the accumulation of nitrogen in agricultural ecosystems and as feed for livestock, particularly in areas where cowpeas are grown [1, 2]. Cowpea is crucial for preventing protein-calorie malnutrition, an excellent supply of vital amino acids (Lys and His), fiber, iron and zinc, as well as significant amounts of bioactive chemicals [3–5]. Aside from these, the protein-rich cowpea seed has additional health advantages, such as a decreased risk of developing cancer and cardiovascular disease. In order to prevent malnutrition caused by a lack of protein and energy in economically challenged areas of developing countries, cowpea provides a high-quality protein component of the daily diet [6, 7]. However, we repeatedly noticed that substantial losses in cowpea yield and production occurred all over the world as a result of various biotic and abiotic challenges such as infections, droughts and salinity [8, 9]. To protect cowpea from these challenges and increase productivity, there is a pressing need.

A wide range of biological processes, including growth and development, epigenetic modification and responses, heterochromatin formation and development patterning are regulated by RNA interference (RNAi) or RNA silencing, which protects plants and other multicellular organisms from biotic and abiotic stressors [10–12]. This defensive mechanism was effective due to transcriptional gene silencing (TGS) and posttranscriptional gene silencing (PTGS). MicroRNA (miRNA) and short interfering RNA (siRNA) are two small RNAs (sRNAs) with a length of 21–24 nucleotides that are involved in both TGS and PTGS. These sRNAs assist plants in building up their natural defenses and protecting themselves against stresses [13–15]. The DCL, AGO, and RDR genes produce the RNAi (DCL, AGO, and RDR) proteins, which are essential for the biosynthesis and activity of sRNAs [16–20]. In this biosynthesis passage, RNA silencing begins with the creation of double-stranded RNAs (dsRNAs), which are then cut into sRNAs that are 21–24 nucleotides long by DCL proteins of the RNase III type [21, 22]. In the reference plant *Arabidopsis*, the miRNA produced by *AtDCL1*, repeat-associated siRNA and the trans-acting siRNA produced by *AtDCL2*, *AtDCL3*, and *AtDCL4* have been linked to virus defense [23]; the strands of sRNA are split after being taken up by the multi-component RNA-induced silencing complex (RISC) [24]. One of the separated strands connected to the AGO proteins serve as a silencing target guide [25]. In some plant species, AGO proteins also regulate transgene silencing [26], epigenetic silencing mechanisms [27], plant development [28] and meristem maintenance [29]. RNA dependent RNA polymerases (RDRs), in addition to DCL and AGO proteins, also produce sRNA. The targeted RNA is amplified to make more secondary dsRNAs, which are then processed into primary siRNAs to repeat the silencing process [13, 30]. As a result, gene engineering can be used to develop future, robust crops that are resistant to stress (biotic and abiotic) by implementing RNA interference technology.

RNAi-related gene families (DCL, AGO, and RDR) have been discovered for a variety of plants and crops, including Brassica, rice, maize, tomato, foxtail millet, pepper, cucumber, sweet orange and others, with respect to AtRNAi genes [31–38]. However, there is a lack of comprehensive knowledge regarding these gene families in cowpea. Therefore, in this study, an effort is made to accomplish a thorough *in silico* analysis for genome-wide identification and characterization of AGO, DCL, and RDR gene families and their associated regulatory elements in cowpea. In Fig. 1, an overview of this study’s workflow in pictures was presented.

**Fig 1.**
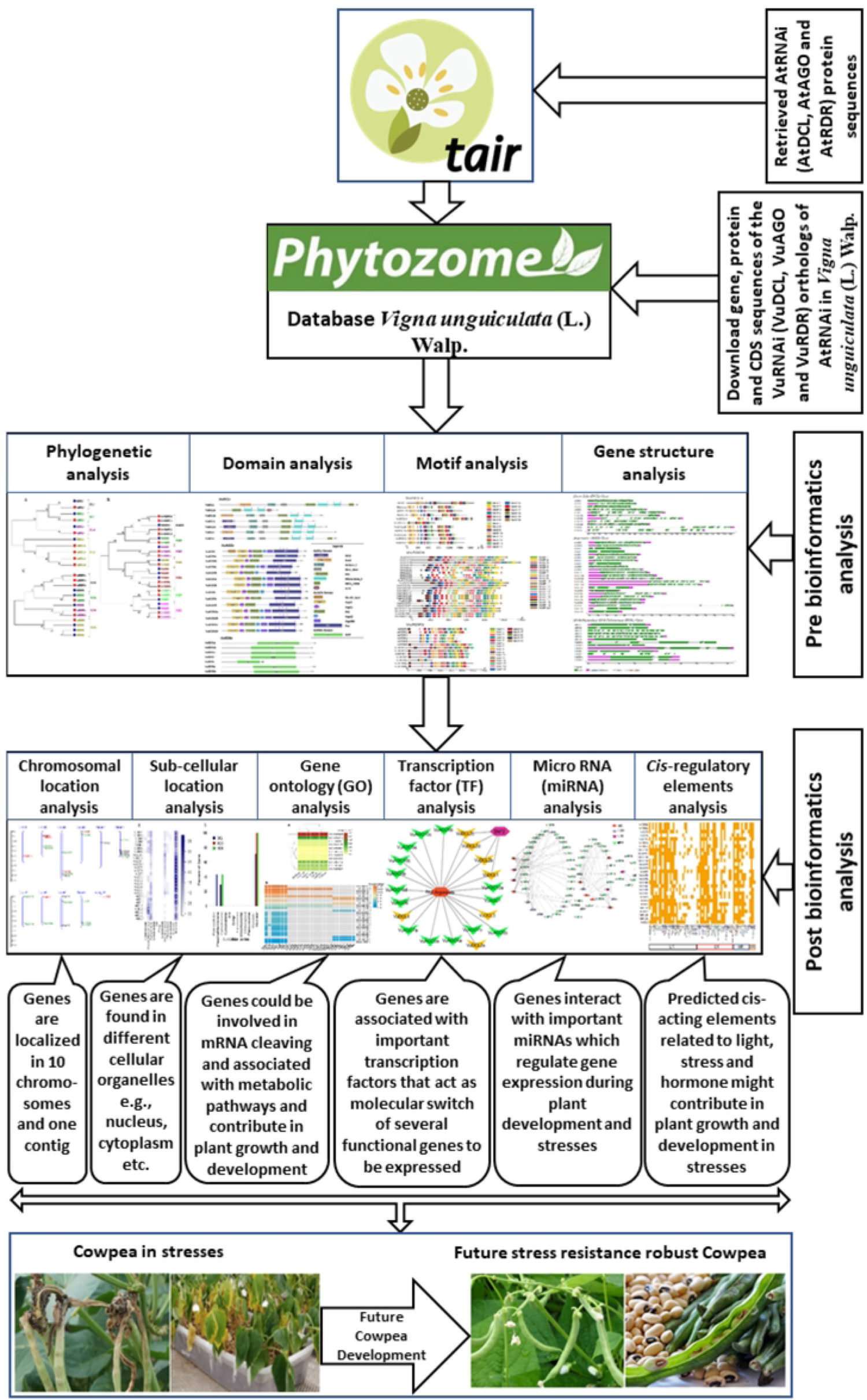
A brief pictorial workflow of this study.

## 2 Materials and Methods

The comprehensive integrated bioinformatics analysis decomposed into three sections 1) discovery of VuRNAi genes 2) characterization of VuRNAi genes and 3) regulatory elements analysis of VuRNAi genes in cowpea.

### 2.1 The data source and descriptions

In order to investigate the cowpea (*Vigna unguiculata*) genome’s VuRNAi genes (DCL, AGO, and RDR), we used its genome and proteome sequences from the Phytozome database (https://phytozome.jgi.doe.gov/pz/portal.html) with Phytozome genome ID: 540, NCBI taxonomy ID: 3917, and website: https://phytozome-next.jgi.doe.gov/info/Vunguiculata v1 2 [39]. Lonardi and his colleagues generated the cowpea genome [40]. By using the query sequences of the AtRNAi genes, this study explored the VuRNAi genes using BLASTP [41] search. 4 AtDCL, 10 AtAGO, and 6 AtRDR sequences totaling 20 AtRNAi genes/proteins were retrieved from the Arabidopsis Information Resource (TAIR) database [42] weblink: (https://www.arabidopsis.org/).

### 2.2 Integrated bioinformatics analyses

The comprehensive bioinformatics analyses comprised BLASTP search, multiple sequence alignment (MSA), phylogenetic tree modeling, functional domain analysis, exon-intron makeup of the RNAi target genes, subcellular localization, GO, TFs, CAREs and miRNA analysis (Fig. 1).

#### 2.2.1 Discovery of VuRNAi genes with respect to AtRNAi genes Exploring VuRNAi genes by BLASTP search

We employed the basic local alignment search algorithm (BLASTP) [41] for proteins to extract the genes, transcripts, CDS and protein sequences for the VuRNAi genes. These VuRNAi gene sequences were retrieved from the Phytozome database [39] using the query sequences AtRNAi protein sequences with an alignment score of 50 and an identity of 50% [38, 43]. To prevent sequence duplication, we solely took into account the primary sequence in the study. The Phytozome database was used to gather the genomic length, protein ID, CDS length, and encoded protein length of the VuRNAi genes. Using the ExPASy database, the molecular weight and isoelectric point (pI) of the pertinent protein sequences were determined [44]. In the supplementary files S1–S3, the full length aligned protein sequences of VuRNAi genes (*VuDCL*s, *VuAGO*s, and *VuRDR*s) were supplied.

### Phylogenetic tree analysis to fix the names of VuRNAi genes

The multiple sequence alignments of the downloaded protein sequences of the candidate the VuRNAi and AtRNAi genes was done to construct phylogenetic tree. This analysis was done using the MEGA7 [45] software. Clustal-W method along with the Neighbor joining method were applied to see the evolutionary relationship in the phylogenetic tree analysis. The best candidate VuRNAi gene were identified comparing the phylogenetic relationship between the considered VuRNAi genes and AtRNAi genes in this analysis.

#### 2.2.2 Characterization of VuRNAi genes

### Analysis of the VuRNAi proteins’ conserved domains and motifs

We used the Pfam database (http://pfam.sanger.ac.uk/) [46] and Multiple Expectation Maximization for Motif Elicitation (MEME-Suite) (https://meme-suite.org/meme/tools/meme) [47] online platforms to investigate the conserved functional domains and motifs of the anticipated VuRNAi protein family. Most major functional domains of VuRNAi proteins that are similar to AtRNAi proteins were kept in the case of conserved functional domain prediction. For the VuRNAi proteins, however, only 20 motifs were taken into account.

### Analysis of VuRNAi gene structures

In this section gene structure of VuRNAi and AtRNAi genes were analyzed to observed the structural similarity between VuRNAi genes with the AtRNAi genes. Comparing the cowpea VuRNAi gene family with its ortholog AtRNAi in *Arabidopsis*, the gene structure of the cowpea gene family was anticipated using the online Gene Structure Display Server (GSDS 2.0, http://gsds.cbi.pku.edu.cn/index.php) [48].

### Localization of VuRNAi genes in chromosomes

Genomic location of the identified *VuDCL*, *VuAGO* and *VuRDR* genes were predicted using online database MapGene2Chromosome V2 (http://mg2c.iask.in/ mg2c_v2.0/).

### Sub-cellular localization of VuRNAi proteins and GO enrichment analysis

The sub-cellular localization of certain proteins controls how the plant cell performs biological tasks. The VuRNAi proteins were located in the cell using a web application called the Plant Subcellular Localization Integrative Predictor (PSI) [49], which may be found at http://bis.zju.edu.cn/psi. Then, GO enrichment analysis was performed using the web tool agriGO v2.0 (http://systemsbiology.cau.edu.cn/agriGOv2/index.php#) [50] to determine if the predicted VuRNAi-related genes were involved in the cluster of different biological processes and molecular functional pathways.

#### 2.2.3 VuRNAi gene regulatory network analysis Regulatory network with transcription factors (TFs)

In this part, we used PlantTFcat: https://www.zhaolab.org/PlantTFcat/, a popular database for plant transcription factor analysis, to examine the regulatory link between the TF family and the anticipated VuRNAi genes. Using Cytoscape 3.7.1, a sub-network of TFs connected with vuRNAi genes was built and visualized in order to identify significant hub TFs and related hub proteins through the interaction network.

### Regulatory network with micro-RNAs

The 19–24 nucleotide long single-stranded non-coding RNA molecules known as microRNAs (miRNAs) are produced by miR genes found in both plants and mammals. Plant growth, development, and stress response are regulated by miRNAs at the transcriptional and post-transcriptional levels. Using mature miRNA sequences and mature miRNA expression in cowpea, we examined the relationship between miRNA and VuRNAi genes in this section using Plant miRNA ENcyclopedia (PmiREN): https://www.pmiren.com/download. Mature miRNA sequence used for target miRNA in plant small RNA using the webserver psRNATarget: https://www.zhaolab.org/psRNATarget/target. Finally, Cytoscape 3.7.1 was used to visualize how the miR genes and VuRNAi genes interact.

### *Cis*-acting regulatory element analysis

The upstream region (1.5 kb of genomic sequences) of each RNAi gene’s start codon (ATG) was excised to study *cis*-acting components of the VuRNAi gene family. Then, using the online prediction analysis tool (PlantCARE): https://bioinformatics.psb.ugent.be/webtools/plantcare/html/ database [51] we predicted the corresponding promoter *cis*-acting regulatory elements. The five categories of LR, SR, HR, OT, and unknown functions were used to categorize the discovered promoter cis-acting regulatory elements.

## 3 Results and discussions

### 3.1 Discovery of VuRNAi gene families in cowpea with respect to AtRNAi genes

To discover VuRNAi genes in the cowpea genome, the *AtDCL*, *AtAGO* and *AtRDR* protein sequences were used as query sequences in a BLASTP search based on the Hidden Markov Model (HMM). In the cowpea genome, we discovered 7 *VuDCL*, 14 *VuAGO*, and 7 *VuRDR* genes. With the neighbor-joining method, phylogenetic trees were generated to name the discovered VuRNAi genes based on how similar they were to the AtRNAi genes (Fig. 2). In the Fig 2 (A) all the predicted 7 *VuDCL*s are clustered into four clades which were differentiated with identical colors and named as DCL1, DCL2, DCL3 and DCL4 according to the number of *AtDCL*s comprising in the clades. The *VuDCL1*, *VuDCL2* (a, b, c, d), *VuDCL3* and *VuDCL4* were clustered with *AtDCL1*, AtDCL2, *AtDCL3* and *AtDCL4* in the clades DCL1, DCL2, DCL3 and DCL4 respectively conforming well-supported bootstrap values. The DCL members are crucial to the sRNA biogenesis process since they help transform dsRNAs into mature sRNAs [52, 53]. *VuDCL1* and *AtDCL1* share a common clade; therefore, it makes sense that their roles would be similar. In light of this, it is likely that *VuDCL1* has a role in development, environmental stress conditions, and flowering mechanisms [54–57]. Accordingly, we hypothesize that *VuDCL2* (a, b, c, d), and *VuDCL3* might regenerate siRNAs and trans-acting small interfering RNA (ta-siRNAs) in accordance with the function of *AtDCL2* and *AtDCL3*, which may be involved in vegetative phase development, disease resistance, and flowering mechanisms [57, 58]. Based on *AtDCL4* function, we may infer that *VuDCL4* may be involved in ta-siRNA metabolism and affect epigenetic maintenance during post-transcriptional silencing through RNA-dependent methylation (RdDM) [23, 59].

**Fig 2.**
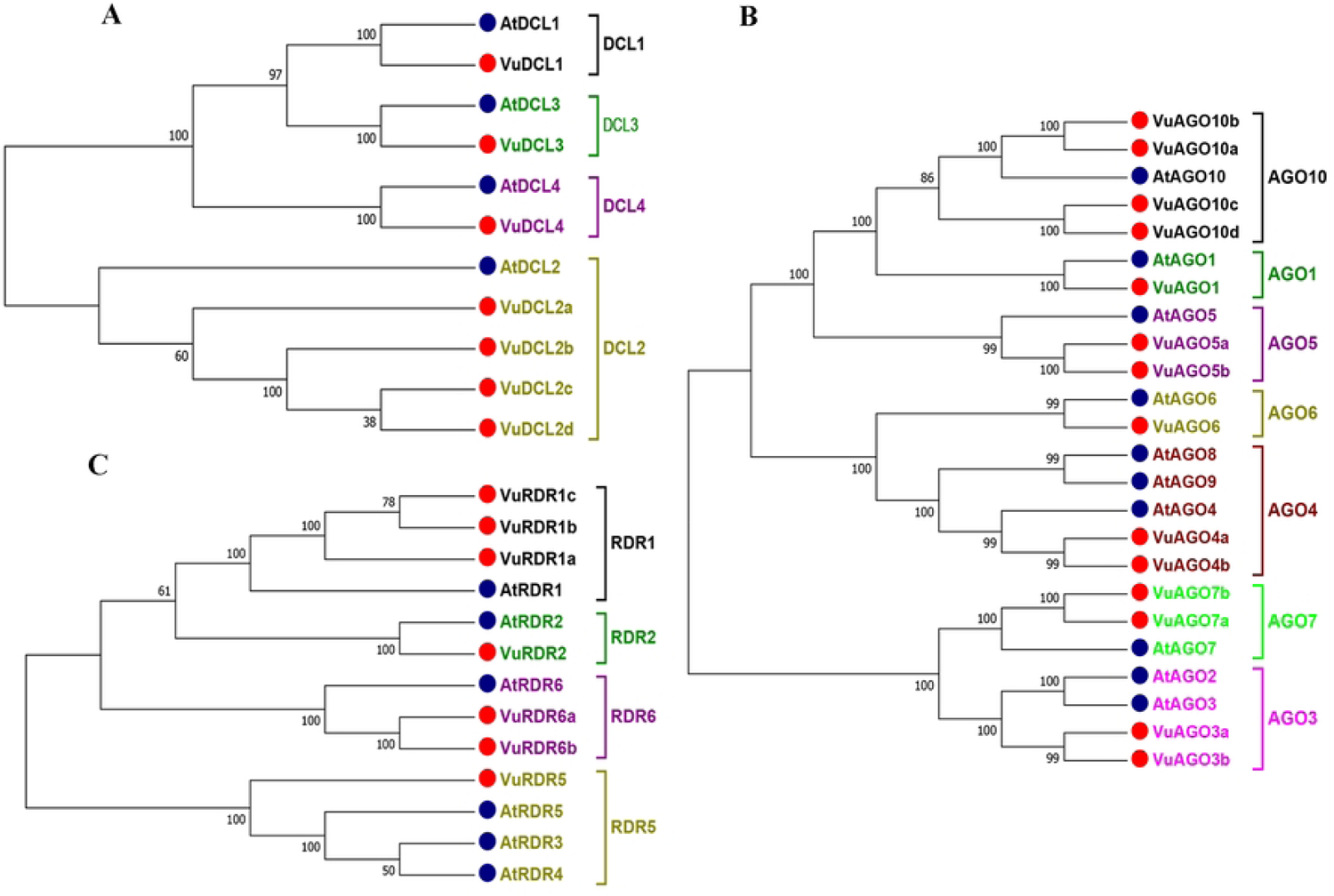
Phylogenetic tree of (A) DCL, (B) AGO and (C) RDR genes in cowpea and their orthologous in *Arabidopsis*. All the trees were constructed using the neighbor-joining method and the on the nodes indicate the percentages of bootstrap values from 1000 replications. In the phylogenetic trees the clades are differentiated using different colors, the red circles represent the VuRNAi genes in cowpea and the blue circles represent their orthologous genes in *Arabidopsis*.

According to Fig. 2 (B), 14 *VuAGO*s were grouped into seven clades based on their highest degree of sequence similarity to *AtAGO*s. The clades were differentiated with identical colors and named as AGO1, AGO3, AGO4, AGO5, AGO5, AGO7 and AGO10. In the figure, *VuAGO1* having the sequence similarity with *AtAGO1* and clustered in clade AGO1. Similarly, *VuAGO3* (a, b) having the highest sequence similarity with *AtAGO3* and then *AtAGO2* and they clustered in clade AGO3. In the same way, *VuAGO4* (a, b) clustered with *AtAGO4*, *AtAGO9* and *AtAGO8* in the clade AGO4, *VuAGO5* (a, b) clustered in the clade AGO5 with *AtAGO5*, *VuAGO6* clustered with *AtAGO6* in the clade AGO6, *VuAGO7* (a, b) clustered with *AtAGO7* in the clade AGO7 and *VuAGO10* (a, b, c, d) clustered with AtAGO10 in the clade AGO10. Ten AtAGO genes in Arabidopsis produce proteins that are important to the RNA silencing mechanism [26, 60]. The role of *AtAGO1* suggests that VuAGO1 may be associated with miRNA and transgene silencing mechanisms [26, 61]. We can predict that *VuAGO4* (a, b) may be involved with endogenous siRNA activity and necessary for epigenetic silencing based on the role of *AtAGO4* [27, 62]. We can anticipate that *VuAGO7* and *VuAGO10* may be necessary for the conversion of plants from the juvenile stage to the adult phase and the development of meristem tissue based on the function of *AtGAO7* and *AtAGO10* [28,29,63].

Based on the phylogenetic tree (Fig 2(C)) there were four clades, *VuRDR (a, b, c)* had their sequence similarity with *AtRDR1* and make a subfamily in the clade RDR1. *AtRDR2* and *VuRDR2* make another clade RDR2 since they have sequence similarity. Similarly, *VuRDR6* (a, b) find their sequence similarity with *AtRDR6* and grouped into the clade RDR6. Finally, *VuRDR5* and *AtRDR5* are same according to the sequence. From the literature it is found that RDR proteins generate dsRNAs from the sRNA and initiate RNA silencing mechanism [64, 65]. We could predict that VuRDR1 is a key component of the RNA silencing pathway and that it might be stimulated by viral infection, salicylic acid, might be involved in antiviral defense and transgene silencing in many species of plants, similar to how *AtRDR1* functions [66–69]. We can infer that *VuRDR2* may contribute to the production of siRNA and be connected to chromatin modification based on the way *AtRDR2* functions [70, 71]. The *AtRDR6* function predicts that *VuRDR6* could generate ta-siRNA precursor and aid in antiviral defense by degrading RNA molecules [72]. These findings demonstrate their functional diversity, which is important for the introduction of strong, stress-resistant cowpea in the future.

Table 1 showed the predicted properties of the VuRNAi genes, such as their location on the chromosome, their structure (ORF length, gene length, and number of introns), and their protein profile (molecular weight of the encoded protein and isoelectric point, or pI). DEAD, Helicase-C, Dicer-dimer, PAZ, RNase III, and DSRM were predicted in the polypeptide sequence of the seven *VuDCL* loci, and they resemble the *AtDCL* all conserved domains (Fig 3). The putative VuDCL genes ranged in genomic length from 8460 base pairs (bp) (*VuDCL2c*: Vigun06g138900) to 24829 bp (*VuDCL3*: Vigun09g219100), with 1218 and 1667 amino acids (aa) of protein coding potential, respectively. Their ORF length, however, ranged from 3654 bp to 5001 bp (Table 1). The pI values of the *VuDCL*s range from 5.87 (*VuDCL4*) to 7.06 (*VuDCL3*), demonstrating the acidic nature of the *VuDCL* genes.

**Fig 3.**
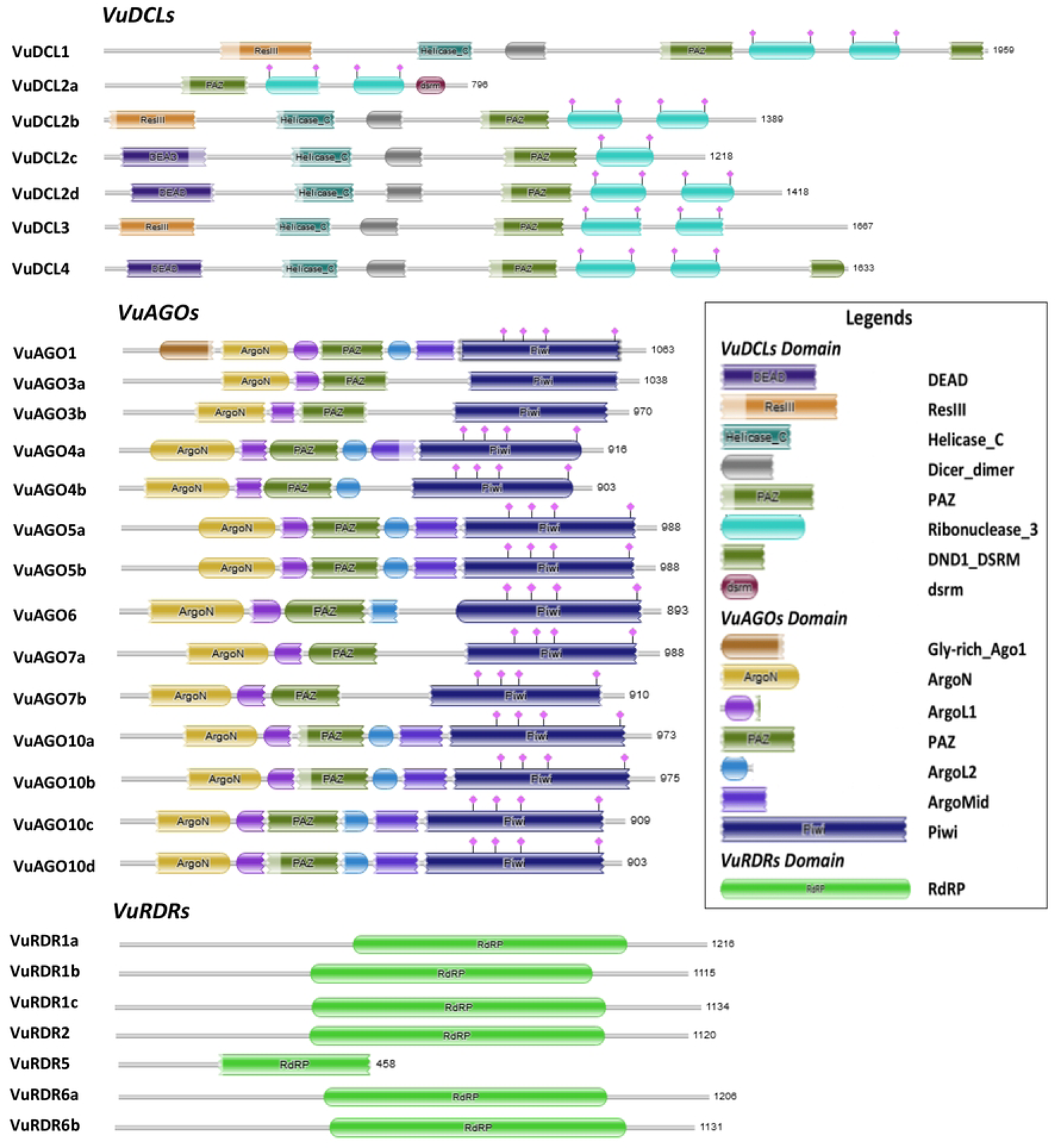
Structure of the conserved domain of the predicted VuDCL, VuAGO and VuRDR proteins analyzed using Pfam database. In the figure predicted domains are indicated by color boxes and at the end of each domains the length of the proteins aa (amino acid) are given.

**Table 1.** Basic properties of the VuRNAi gene families in cowpea.

| Gene properties |  |  |  |  |  |  | Protein properties |  |  |
| --- | --- | --- | --- | --- | --- | --- | --- | --- | --- |
| Serial # | Gene name | Accession ID | Chromosomal location | ORF length (bp) | Gene length (bp) | Number of intron | Molecular weight (KD) | Protein length (aa) | pI |
| VuDCL |  |  |  |  |  |  |  |  |  |
| 1 | *VuDCL1* | Vigun09g003800.1 | Vu09:275033-288198 | 5877 | 13165 | 19 | 219.49 | 1959 | 6.28 |
| 2 | *VuDCL2a* | Vigun06g122700.1 | Vu06:25020658-25032899 | 2388 | 12241 | 8 | 90.85 | 796 | 7.02 |
| 3 | *VuDCL2b* | Vigun06g139200.1 | Vu06:26462778-26478925 | 4167 | 16147 | 21 | 157.20 | 1389 | 7.05 |
| 4 | *VuDCL2c* | Vigun06g138900.1 | Vu06:26440174-26448634 | 3654 | 8460 | 20 | 138.26 | **1218** | 6.88 |
| 5 | *VuDCL2d* | Vigun05g132600.1 | Vu05:15578574-15589149 | 4254 | 10575 | 22 | 160.38 | 1418 | 6.77 |
| 6 | *VuDCL3* | Vigun09g219100.1 | Vu09:39292521-39317350 | 5001 | 24829 | 24 | 187.79 | 1667 | 7.06 |
| 7 | *VuDCL4* | Vigun03g367100.1 | Vu03:57043142-57063002 | 4899 | 19860 | 24 | 184.29 | 1633 | 5.87 |
| VuAGO |  |  |  |  |  |  |  |  |  |
| 1 | *VuAGO1* | Vigun04g040700.1 | Vu04:3512565-3519416 | 3189 | 6851 | 20 | 117.27 | 1063 | 9.38 |
| 2 | *VuAGO3a* | Vigun06g143400.1 | Vu06:26898561-26903499 | 3117 | 4938 | 2 | 117.03 | 1039 | 9.01 |
| 3 | *VuAGO3b* | Vigun03g198800.1 | Vu03:28537393-28540930 | 2913 | 3537 | 2 | 109.05 | 971 | 9.03 |
| 4 | *VuAGO4a* | Vigun06g025400.1 | Vu06:11767096-11775775 | 2748 | 8679 | 21 | 102.40 | 916 | 8.80 |
| 5 | *VuAGO4b* | Vigun08g178900.1 | Vu08:34878837-34886097 | 2709 | 7260 | 21 | 100.81 | 903 | 9.14 |
| 6 | *VuAGO5a* | Vigun11g132800.1 | Vu11:34148188-34154497 | 2964 | 6309 | 21 | 109.30 | 988 | 9.66 |
| 7 | *VuAGO5b* | Vigun11g133000.1 | Vu11:34188493-34194747 | 2964 | 6254 | 21 | 109.13 | 988 | 9.66 |
| 8 | *VuAGO6* | Vigun11g043400.1 | Vu11:6379474-6395631 | 2679 | 16157 | 2 | 100.04 | 893 | 8.76 |
| 9 | *VuAGO7a* | Vigun02g055700.1 | Vu02:19910434-19915778 | 2967 | 5344 | 2 | 112.57 | 989 | 9.28 |
| 10 | *VuAGO7b* | Vigun02g055800.1 | Vu02:19950731-19954473 | 2730 | 3742 | 1 | 103.24 | 910 | 9.25 |
| 11 | *VuAGO10a* | Vigun07g003200.1 | Vu07:253781-261378 | 2919 | 7597 | 20 | 108.95 | 973 | 9.26 |
| 12 | *VuAGO10b* | Vigun07g233900.1 | Vu07:35574238-35582483 | 2925 | 8245 | 20 | 109.60 | 975 | 9.27 |
| 13 | *VuAGO10c* | Vigun09g063300.1 | Vu09:6634205-6647708 | 2727 | 13503 | 21 | 103.27 | 909 | 9.10 |
| 14 | *VuAGO10d* | Vigun03g350900.1 | Vu03:55202472-55215146 | 2709 | 12674 | 21 | 102.16 | 903 | 9.05 |
| VuRDR |  |  |  |  |  |  |  |  |  |
| 1 | *VuRDR1a* | Vigun02g017800.1 | Vu02:6314155-6322447 | 3651 | 8292 | 5 | 138.87 | 1217 | 8.58 |
| 2 | *VuRDR1b* | Vigun08g009200.1 | Vu08:818516-822196 | 3345 | 3680 | 4 | 126.55 | 1115 | 6.44 |
| 3 | *VuRDR1c* | Vigun02g017700.1 | Vu02:6301580-6310988 | 3402 | 9408 | 3 | 129.40 | 1134 | 8.28 |
| 4 | *VuRDR2* | Vigun03g326600.1 | Vu03:52257972-52262538 | 3360 | 4566 | 3 | 127.39 | 1120 | 6.23 |
| 5 | *VuRDR5* | Vigun04g008600.1 | Vu04:614376-620001 | 1377 | 5625 | 9 | 51.78 | 459 | 8.20 |
| 6 | *VuRDR6a* | Vigun01g151600.1 | Vu01:33427336-33433259 | 3618 | 5923 | 1 | 137.47 | 1206 | 7.97 |
| 7 | *VuRDR6b* | VigunL057500.1 | contig_465:34900-40824 | 3393 | 5924 | 1 | 128.83 | 1131 | 8.17 |

On the other hand, the polypeptide sequences of the 14 *VuAGO* genes, which are the primary members of the plant AGO protein family, were shown to include the most significant conserved domains, such as an N-terminal PAZ and a C-terminal PIWI, based on the HMM analysis (Fig 3). The found *VuAGO* genes have genomic lengths ranging from 3537 base pairs (bp) (*VuAGO3b*: Vigun03g198800) to 16157 bp (*VuAGO6*: Vigun07g003200), with 971 and 893 amino acids (aa) of potential protein coding potential, respectively. However, they have respective ORF lengths of 2679 bp and 2913 bp (Table 1). *VuAGO*s’ pI values, which ranged from 8.76 (VuAGO6) to 9.66 (*VuAGO5a* and *VuAGO5b*) (Table1).

Our HMM analysis predicted that the *VuRDR* gene family will have the RdRP conserved domain, just like the other plants, including A. thaliana (Fig 3). The seven discovered *VuRDR*s ranged in genomic length from 3680 bp (*VuRDR1b*: Vigun08g009200) to 9408 bp (VuRDR1c: Vigun02g017700), with respective protein coding potentialities of 1115 aa and 1134 aa. Their respective ORF lengths are 3345 bp and 3402 bp. The *VuRDR*s proteins’ pI values, which vary from 6.23 to 8.58, demonstrated that the proteins are more likely to be acidic (Table1).

### 3.2 Conserved domains and motifs analysis of VuRNAi proteins

According to Fig. 3, the functional domains of the DCL, AGO, and RDR protein families are well conserved in the *VuDCL*, *VuAGO*, and *VuRDR* proteins. The majority of the cowpea *VuDCL* proteins contain the important domains DEAD, /Res III, Helicase-C, Dicer-dimer, PAZ, and Ribonuclease-3/RNase III; however, the dsrm domain is only present in *VuDCL2a* (Fig 3). Different plants, including orange, banana, wild sugarcane and *Arabidopsis*, also produced comparable results [38,43,73]. The anticipated domains are crucial for protein function in plants [59, 74]. Plants with two DCL genes (DCL2 and DCL3) that protect them from viral infection [75]. The dsRNAs are split into 21–24 nucleotide long sRNAs by the DCLs proteins. The two Ribonuclease-3/RNase III catalytic functional domains cleave siRNA and dsRNA, which are primarily the tasks of the PAZ domain. These sRNAs provide the endonuclease enzyme-containing RNA-induced silencing complex (RISC), which provokes the AGO proteins to cleave the target homologous RNAs in accordance with the sRNAs’ structural arrangement [32, 55]. The AGO proteins identify two crucial domains: N-terminal PAZ and C-terminal PIWI [76–78]. All of the *VuAGO* proteins were expected to contain these domains (PAZ and PIWI). These outcomes have also been reported for *AtAGO* proteins (AGO functional domains)[79]. Additionally, it is stated in the literature that the PAZ and PIWI domains of AGOs are crucial for RNase activity [77, 80]. Both domains shared the same homology with RNase H, which binds to the 5’ end of the siRNA of the target RNA and cleaves it, proving the complementary nature of the sRNAs [81, 82]. The *VuAGO* proteins’ projected PAZ and PIWI conserved domains may have a crucial functional role in converting double-stranded RNA into single-stranded RNA and in stimulating the target RNA degradation process [80, 81]. Similar to *AtAGO1*, which coordinates ribosome binding to promote AGO protein stimulation for the RNA silencing process, the Gly-rich Ago1 domain predicted in the *VuAGO1* protein stimulates RNA silencing [83]. By creating dsRNAs utilizing single-stranded RNAs (ssRNAs) as templates, the RDRs proteins assist in initiating a new RNAi silencing process. RDR proteins have a single conserved RdRP domain that contains a catalytic β’ subunit of the RdRP motif [84–86]. Our study demonstrated that all *VuRDR* proteins have the usual RdRP domain, which is identical to the RdRP conserved domain of *AtRDR*s (Fig 3).

The motif analysis of *VuDCL*, *VuAGO* and *VuRDR* proteins using MEME-suite database and TBtools [87] presented in Fig 4. From the figure it is observed that *VuDCL1* has 20 motifs *VuDCL*(*2a, 2b, 2c* and *2d*) have 10, 20, 18 and 20 motifs respectively. *VuDCL3* and *VuDCL4* have 19 and 18 motifs respectively. The results are almost similar to the AtDCL proteins. There are 10 AtAGO proteins ranges from *AtAGO1* to *AtAGO10*. However, the paralogs of *AtAGO2, AtAGO8 and AtAGO9* are absent in cowpea. We considered a maximum of 20 motifs for the VuAGO proteins in the analysis. Among which *VuAGO1* contains 20, each of *VuAGO* (3a and 3b) contains 18, each of *VuAGO* (4a and 4b) contains 17, each of *VuAGO* (5a and 5b) contains 20, each of *VuAGO* (*7a* and *7b*) contains 18, each of *VuAGO* (10a, 10b, 10c and 10d) contains 20 and *VuAGO6* contains 18 motifs (Fig 4). These results exhibit higher conservation with their respective paralogs of *AtAGO* proteins. The MEME-suite analysis of *VuRDR* proteins considering 20 motifs. The *VuRDR* proteins have motifs ranges 17-20 except *VuRDR5* having 4 motifs (Fig 4).

**Fig 4.**
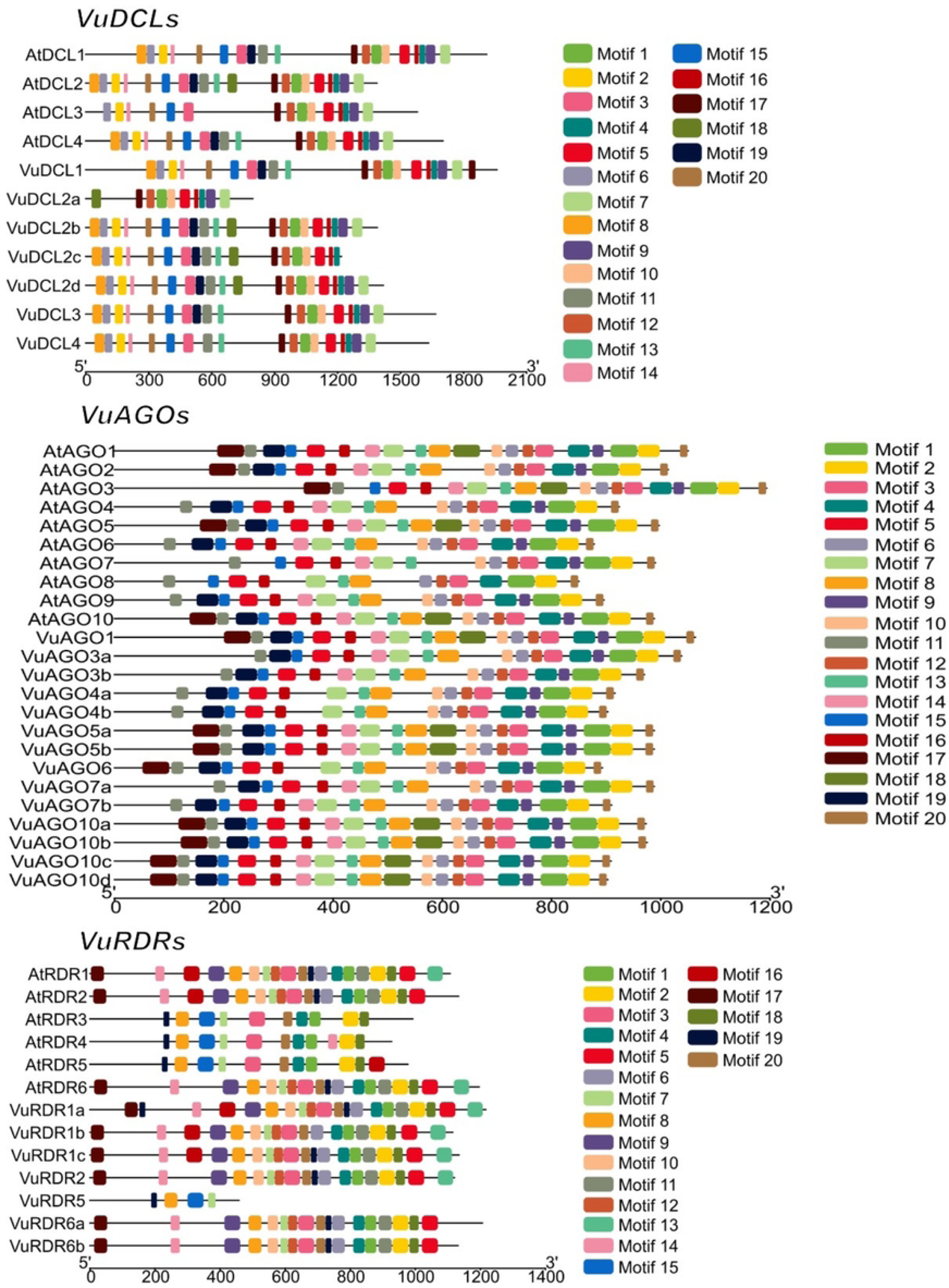
Structure of the conserved motif (maximum of 20 motifs are displayed) of the predicted VuDCL, VuAGO and VuRDR proteins analyzed using MEME-suite. Each color represents different motifs in the predicted proteins and the scale bars presented at the bottom of each plot (VuDCLs, VuAGOs and VuRDRs) represents the length of proteins in aa (amino acid).

### 3.3 VuRNAi genes structures analysis with respect to AtRNAi genes

*VuDCL*s, *VuAGO*s, and *VuRDR*s show well-conserved gene structure similar to AtDCLs, AtAGOs, and AtRDRs of Arabidopsis, according to gene structure analysis utilizing the Gene Structure Display Server (GSDS) (http://gsds.gao-lab.org/) (Fig 5). Exon-intron number (20-19) for *VuDCL1* and *VuDCL2b* were similar same to the *AtDCL1 and AtDCL2* (20-19) and (22-21) respectively. This number for *VuDCL2c/d* is very close to the *AtDCL2* except *VuDCL2a* (9-8). The exon-intron number for the *VuDCL4* (24-23) is also very close to *AtDCL4* (25-24), while it is to some extent different between *VuDCL3* and *AtDCL3*. On the other hand, exon-intron numbers for *VuAGO1*, *VuAGO3a/b*, *VuAGO4a*, *VuAGO6*, *VuAGO7a* are exactly similar with the *AtAGO1*, *AtAGO3*, *AtAGO4*, *AtAGO6* and *AtAGO7* which are 21-20, 3-2, 22-21, 22-21 and 3-2 respectively. The ranges of exon and intron for VuAGO10a/b/c/d are 21-22 and 20-21 respectively that are very similar to the *AtAGO10* (19-18) (Fig 5). Subsequently, exon-intron numbers for the *VuRDR2* and *VuRDR6a/b* are exactly similar to the *AtRDR2* and *AtRDR6* that are 4-3 and 2-1 respectively. The other *VuRDR*s are more or less similar exon-intron number compare to the respective *AtRDR*s (Fig 5). This analysis led us to the conclusion that *VuDCL*, *VuAGO* and *VuRDR* gene structures are more comparable to those of their orthologs (*AtDCL*, *AtAGO* and *AtRDR*) in Arabidopsis, indicating that these genes have functionally similar roles in the RNAi pathway.

**Fig 5.**
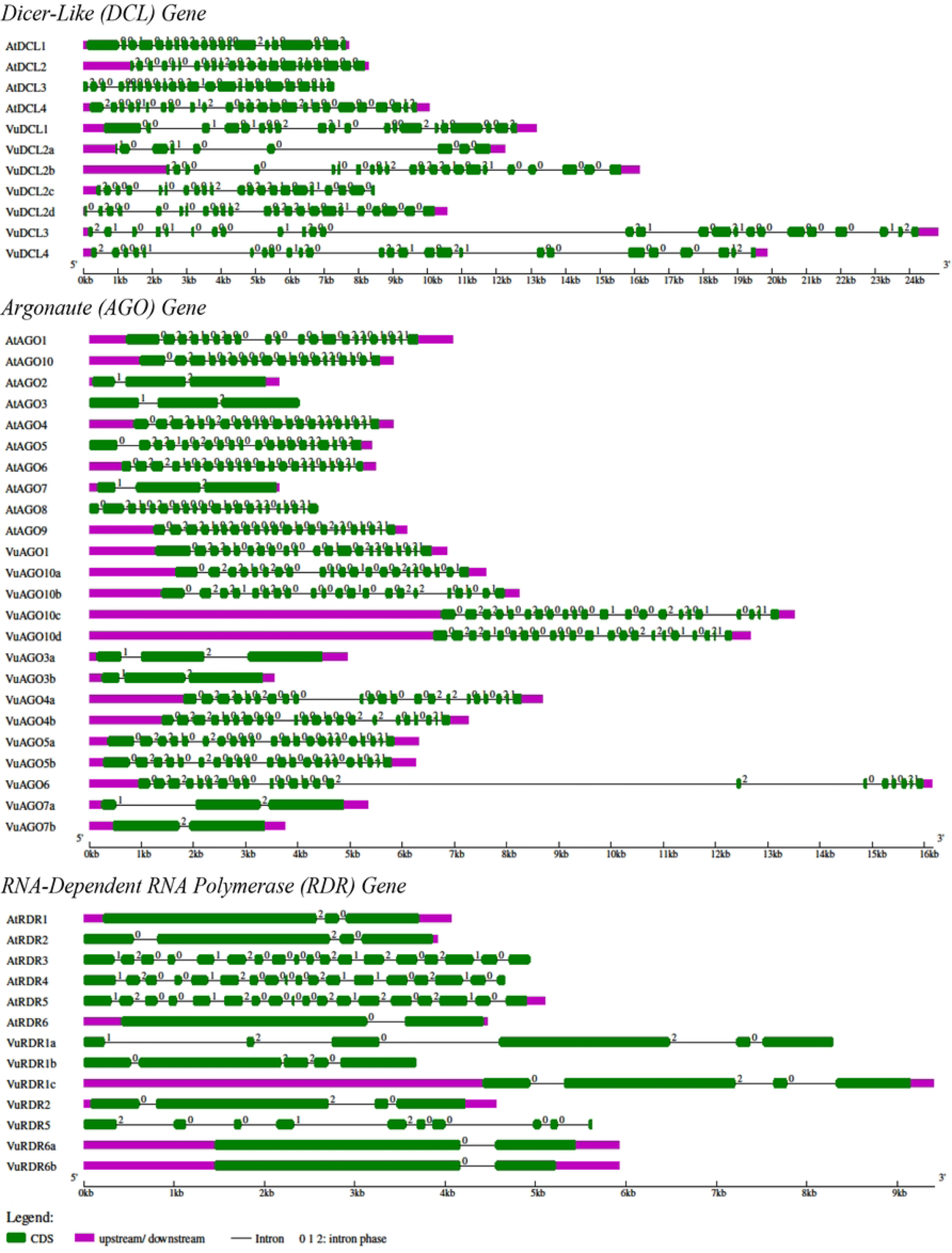
Structural presentation of VuDCL, VuAGO and VuRDR genes along with AtDCL, AtAGO and AtRDR genes.

### 3.4 Localization of VuRNAi genes in chromosomes

The cowpea (*Vigna unguiculata* (L.) Walp.), which has 22 chromosomes and a relatively modest 620 Mb genome, is closely linked to other legume crops [88]. The predicted 28 cowpea VuRNAi genes encoding DCLs, AGOs and RDRs proteins localized on 10 chromosomes and one contig (Fig 6). Among the 7 *VuDCL* genes *VuDCL1 and VuDCL3* are located on the chromosome 09, *VuDCL2a, VuDCL2b, VuDCL2c* were located on chromosome 06 and *VuDCL2d* mapped on chromosome 05 and *VuDCL4* is plotted on the chromosome 03. There are 14 *VuAGO*s among those *VuAGO7a* and *VuAGO7b* were plotted on chromosome 02, *VuAGO3b* and *VuAGO10d* mapped on chromosome 03, *VuAGO1* located on the chromosome 04, *VuAGO3a* and *VuAGO4a* were located on the chromosome 06, *VuAGO10a* and *VuAGO10b* were located on chromosome 07, chromosome 08 contains *VuAGO4b*, chromosome 09 contains *VuAGO10c* and *VuAGO06, VuAGO5a and VuAGO5b* were plotted on the chromosome 11. The distribution of the 7 VuRDRs over 10 chromosomes and 1 contig was that the chromosome 01 contains *VuRDR6a, VuRDR1a* and *VuRDR1c* mapped on the chromosome 02, *VuRDR2 and VuRDR5* were plotted on the chromosome 03 and chromosome 04 respectively, chromosome 08 contains *VuRDR1b* and lastly *VuRDR6b* is located on the contig-465.

**Fig 6.**
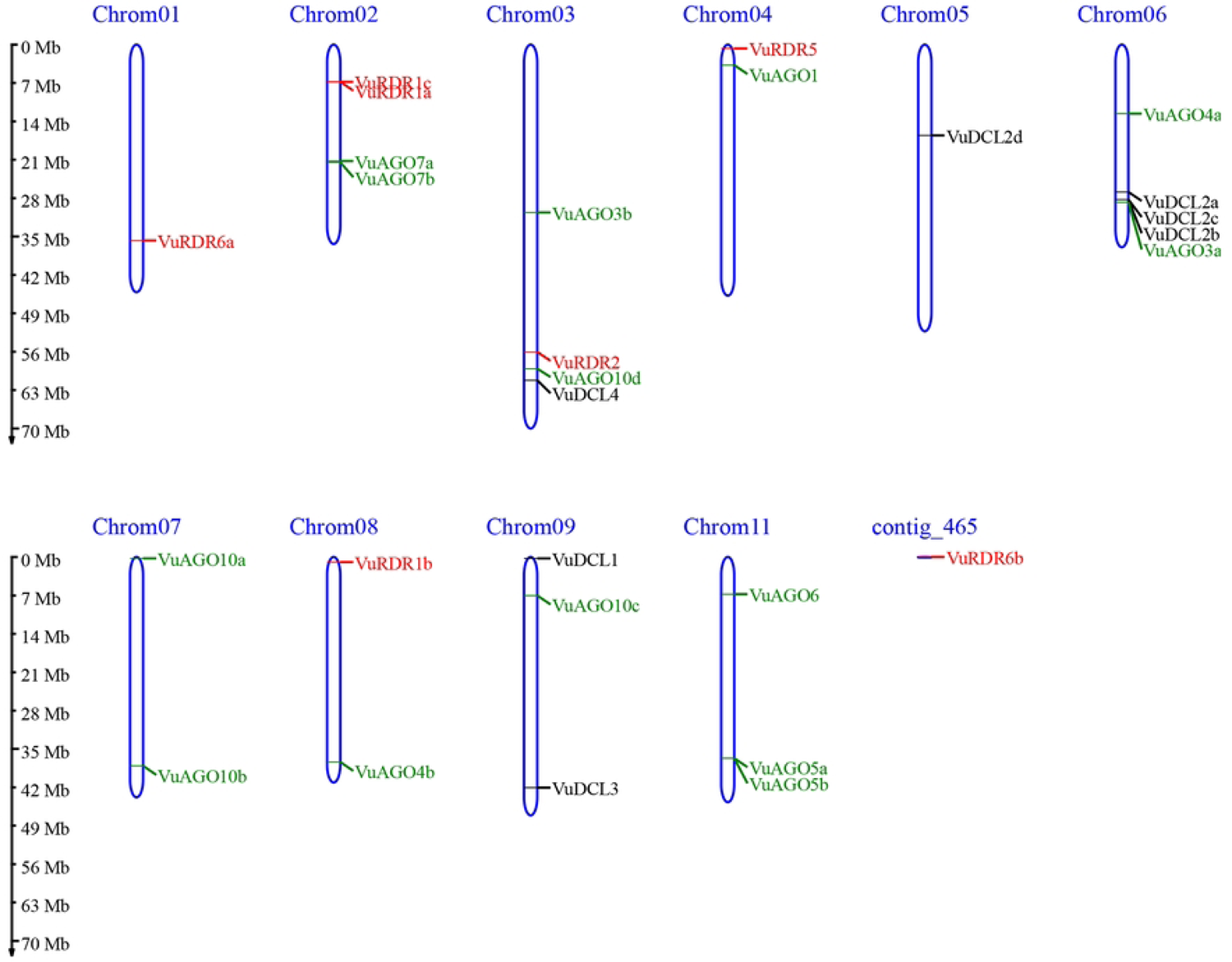
Genomic localization of VuDCL, VuAGO and VuRDR genes. In the figure, the chromosome number and the contig number are shown at the top of the bar (chromosome). In the graph, length of the chromosome shown at the left side scaling in mega base (Mb).

### 3.5 Sub-cellular localization of VuRNAi proteins

There are two types of RNAi-mediated gene silencing: transcriptional gene silencing (TGS) and posttranscriptional gene silencing (PTGS). The TGS take place in the nucleus through DNA methylation and histone modification, on the other hand PTGS completed in the cytoplasm through degrading mRNA [19,89–91]. Therefore, sub-cellular localization of the RNAi proteins regulate expression of targeted genes in eukaryotic cells as well as the functional forms of proteins depends on their location at the cellular level [92, 93]. We observed that *VuDCL*, *VuAGO* and *VuRDR* proteins are localized in the nucleus, plasma membrane, cytoplasm, and mitochondria (Fig 7(A)). Nonetheless, the presence of these proteins in the mitochondria is insignificant (Fig 7(A)). According to this proclamation, *VuDCL1* present in the cytoplasm and nucleus, *VuDCL2a* localize only in the nucleus, VuDCL(2a,2c) located only in the plasma membrane, *VuDCL2d* present in the plasma membrane, cytoplasm, and nucleus, *VuDCL3* and 4 localize only in the nucleus significantly. It’s interesting to note that all of the VuAGO proteins were significantly localized in the nucleus. Two of the six *VuRDR* proteins, *VuRDR1a* and 1b, were significantly present in both the cytoplasm and nucleus. Even though there was a lot of *VuRDR1c* in the nucleus, there was a lot of *VuRDR2* in both the cytoplasm and the nucleus as well. The rest of the *VuRDR* proteins (*VuRDR5, VuRDR6a and VuRDR6b*) present significantly only in the nucleus. All most similar results also find in the literature [38, 43]. From the Fig 7 (B) it is observed that all VuRNAi genes except *VuDCL2b* and *VuDCL2c* significantly present in the nucleus, more than 40% and 25% of the discovered VuRDR and VuDCL genes significantly present in the cytoplasm respectively and only *VuDCL* (40%>) proteins was present in the plasma membrane. From these evidences it can be conclude that due to the positional appearance of the VuRNAi genes, they might be participated in the TGS and PTGS. As a result, the PTGS process’s RISC-mediated cleavage activities directly include the proteins DCL, AGO, and RDR [94]. Lastly, our computer-based prediction gives important clues about what roles the *VuDCL*, *VuAGO*, and *VuRDR* proteins play in the RNAi pathway.

**Fig 7.**
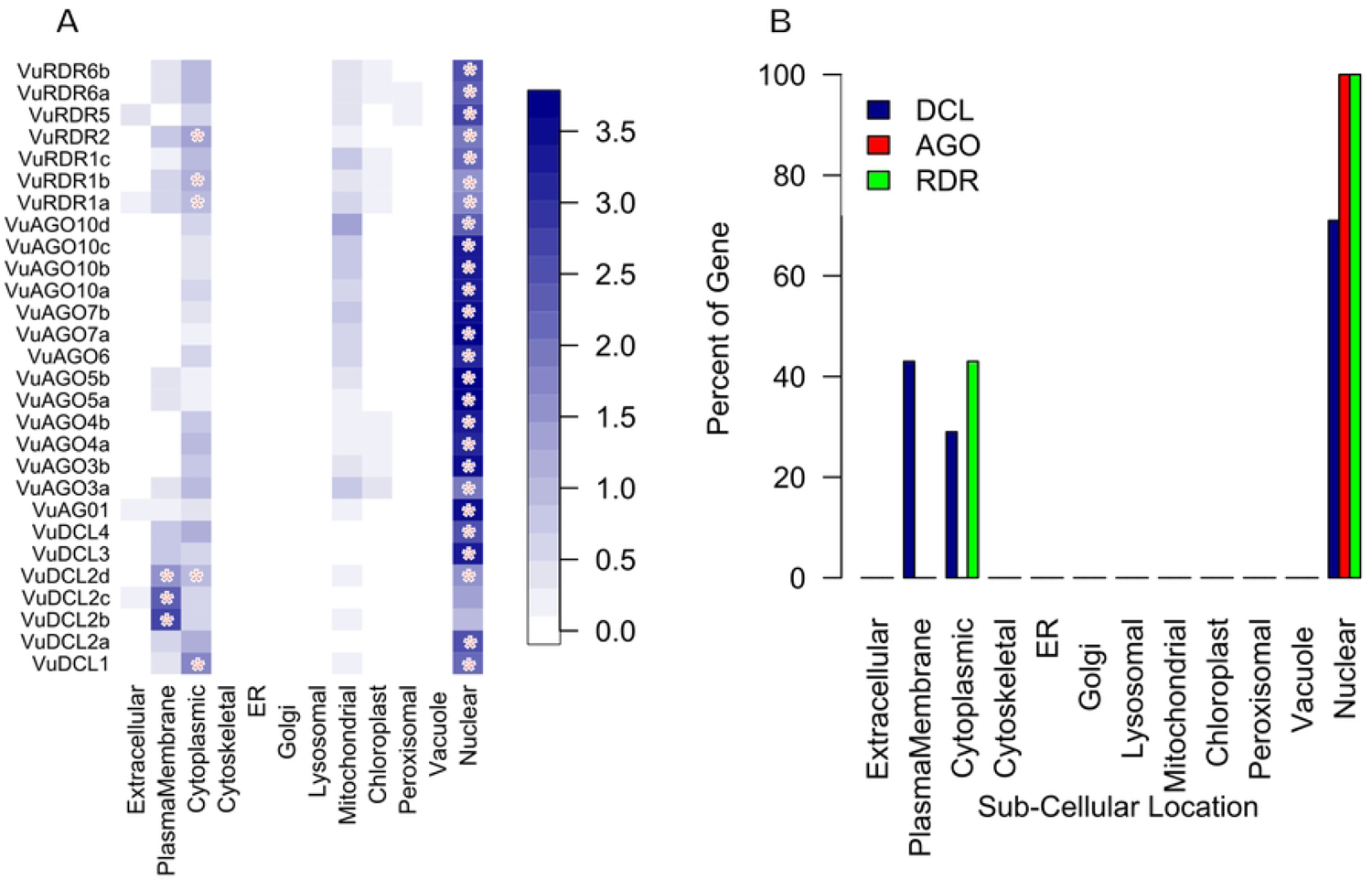
Sub-cellular location analysis for VuDCL, VuAGO and VuRDR proteins. In the figure “A” color deepness represents intensity of presence of the genes in the respective sub-cellular location and the sign “*” represents significant presence of the gene. Figure “B” represents percentage of significant proteins in the sub-cellular location.

### 3.6 Gene ontology (GO) enrichment analysis

The GO enrichment analysis of the VuRNAi proteins was performed to determine the relationship between the VuRNAi proteins and various biological processes and molecular activities. The GO terms associated with the VuRNAi proteins characterize functions of VuRNAi genes in gene silencing (Fig 8 and file S4). According to the results 7 proteins participate in RNA processing (GO:0006396; p-value= 1.70E-06). RNA processing machinery associated with m^6^A-RNA methylation and m^6^A-RNA sites on numerous clock gene transcripts; targeted suppression of m^6^A-RNA methylation by silencing Mettl3 [95]. These 7 VuRNAi proteins are also participate in RNA metabolic process (GO:0016070; p-value=0.0029) and the metabolic process boost the immune system [96]. There are also 7 VuRNAi proteins engaged in nucleobase, nucleoside, nucleotide and nucleic acid metabolic process in cowpea. According to the discussion we can conclude that these proteins are participate in the mRNA degradation in cowpea (Fig 8 (A)).

**Fig 8.**
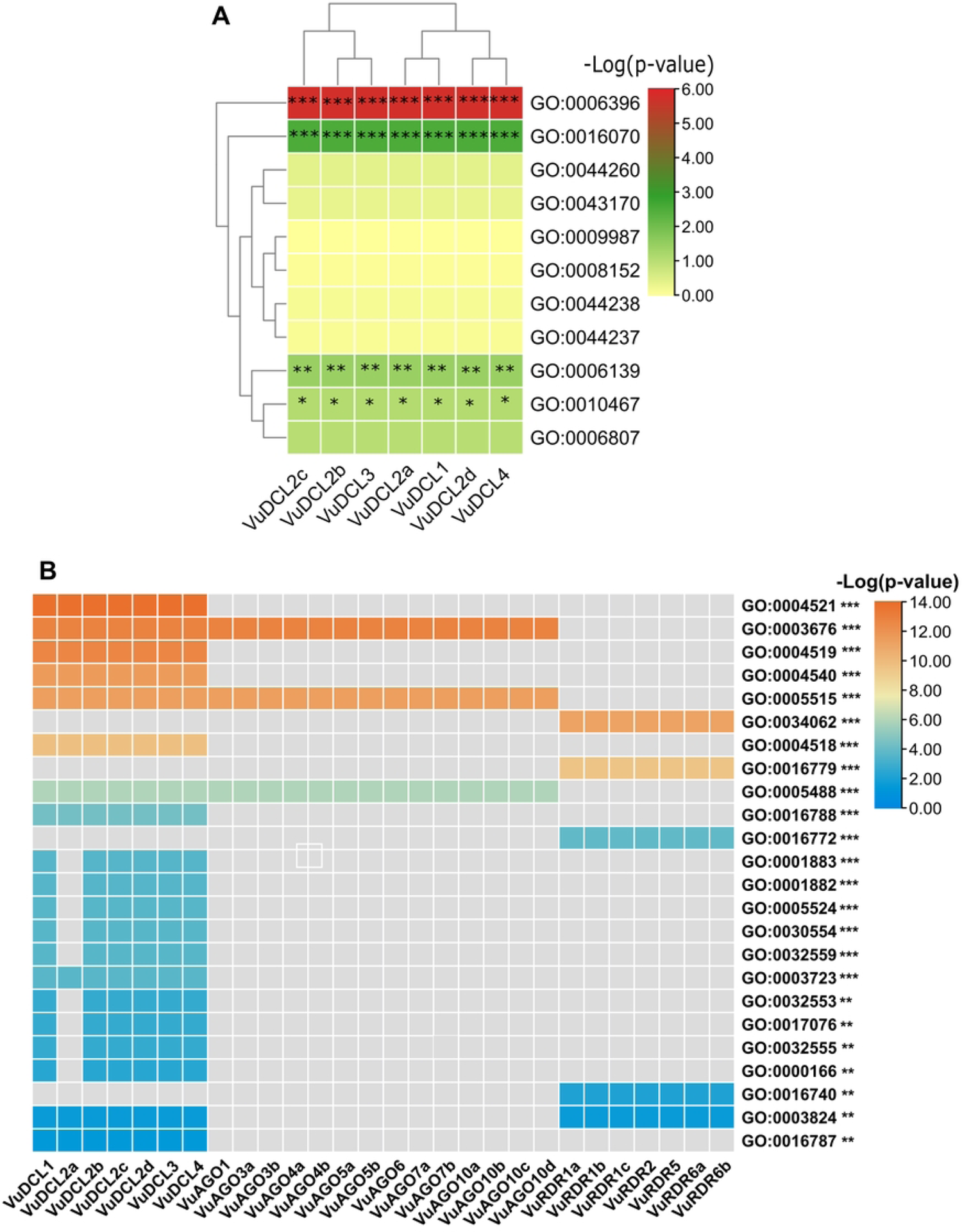
Heatmap of the predicted GO terms and associated VuRNAi gene family. In the figure A) represents biological process and B) represents molecular function. Gray color in the figure B represents absence of the genes in the respective molecular function.

RNAi proteins silence genes through PTGS mechanism in plants and animals [97]. The GO enrichment analysis asserts that VuRNAi genes might participate in the PTGS mechanism through cleaving mRNA in cowpea (Fig 8). Therefore, VuRNAi have close link with RISC which degrades the proteins in cowpea. In this protein degradation process AGO a group of RNAi proteins play the key role by means of endonucleolytic activity in the cell which results PTGS for mRNA substrate [94]. According to figure 8, out of 28 VuRNAi proteins, 7 were engaged in nucleic acid binding activity (p-value = 1.30E-13 for GO:0003676). 7 of the 28 VuRNAi proteins had endonuclease activity (GO:0004519: p-value = 2.20E-13), which breaks internal strands of nucleic acids to form links between them. In addition to breaking internal connections within ribonucleic acid, 7 of the 28 VuRNAi proteins were ribonuclease active (GO:0004540: p-value = 2.60E-12). The protein-binding activity of 21 out of 28 VuRNAi proteins was also detected (GO:0005515, p-value = 3.90E-12). Besides these, the VuRNAi proteins significantly participates in RNA polymerase activity, nuclease activity, ATP binding activity, RNA binding activity, ribonucleotide binding activity etc. Based on the GO enrichment analysis we can conclude that VuRNAi proteins involved in various biological process and functional activity.

### 3.7 Identification of VuRNAi gene regulatory factors

#### 3.7.1 Regulatory network with transcription factors (TFs)

The expression of genes is controlled by transcription factors (TFs), which also have a substantial impact on a number of biological processes in living things, including plants. TFs affect growth, development, metabolism, and defense against microbial invasion. They do this by acting as a molecular switch for many functional genes that are differentially expressed in hair in response to biotic and abiotic stresses [98–101]. These defense mechanism are conducted by the RNAi gene family through their endonucleases activity or by cleaving targeted mRNA and controlling gene expression [102, 103]. In the Fig 9 PAZ-Argonaute family of TF have connection with all the *VuDCL* and *VuAGO* proteins is responsible for endonucleases activity and gene silencing [102, 103]. There is another TF family SNF2 in the same figure is linked with the *VuDCL1, VuDCL2b, VuDCL2c and VuDCL2d* is engaged in gene silencing [104, 105].

**Fig 9.**
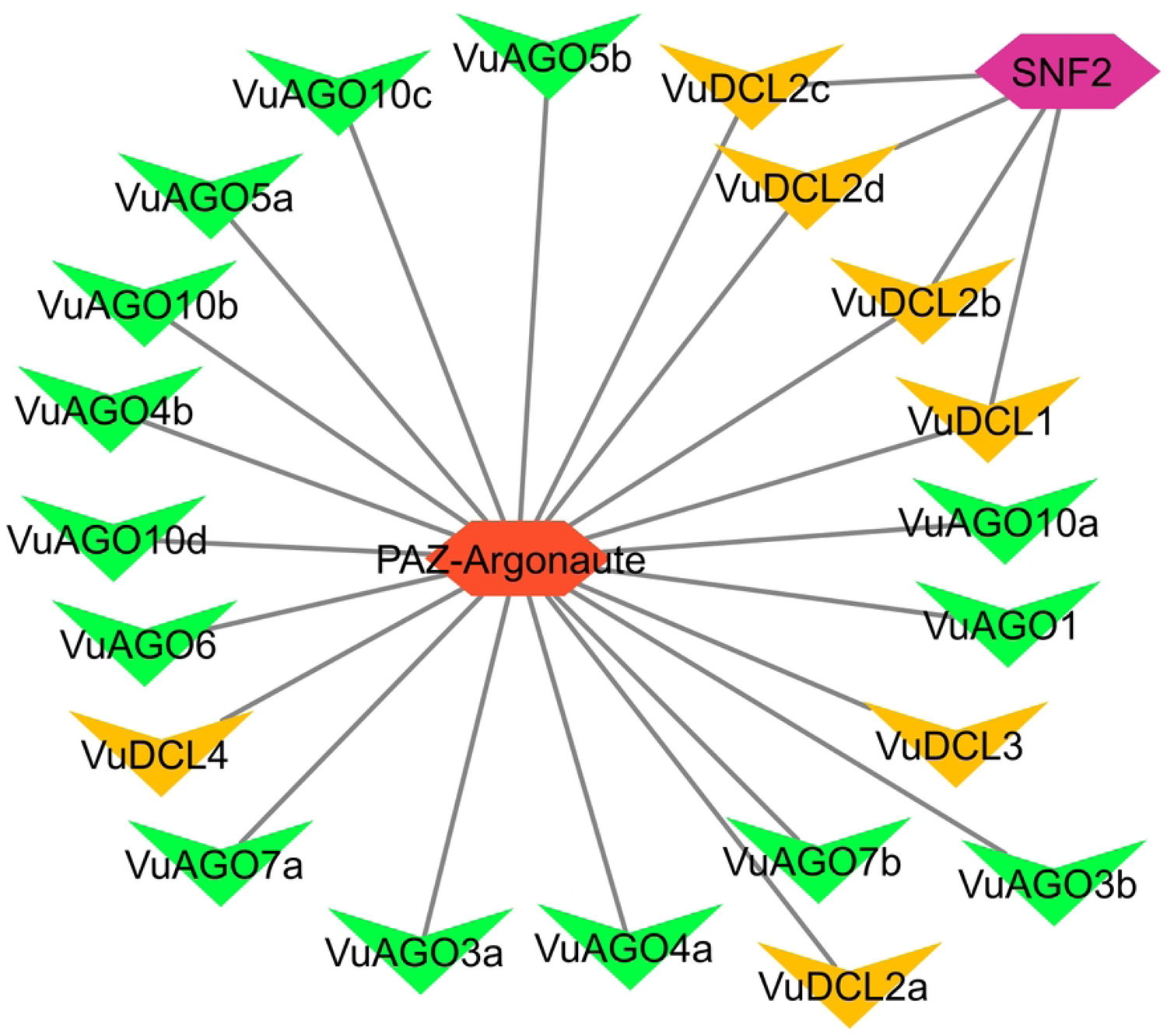
Regulatory network between the VuRNAi genes and TF. In the figure, green and yellow color represent VuAGOs and VuDCLs repectively. Red and purple colors represent TF.

#### 3.7.2 Regulatory network with micro-RNAs (miRNAs)

Development, epigenetic changes, and biotic and abiotic stresses all include the RNA silencing mechanism, which is mediated by the three primary RNAi proteins DCL, AGO, and RDR [14,15,106]. The 19–24 nucleotide single-stranded non-coding RNA molecules known as microRNAs (miRNAs), which have undergone evolutionary conservation, control the expression of certain genes or gene clusters at the TGS and PTGS levels [19,89–91,107]. The miRNAs encoded by *miR* genes generate through the RNAi protein mediated miRNA biogenesis process [106, 108]. In this study, we found 23 miR family genes that work with VuRNAi family genes to synthesize miRNAs. From the Fig 10 we observed that *VuDCL1* linked with 15 miRNAs genes, each of *VuDCL4, VuRDR6a and VuDRD6b* linked with 11 miRNAs genes, *VuDCL2b* linked with 9 miRNAs genes, each of *VuAGO5a and VuAGO5b* linked with 7 miRNAs genes etc. Among these linked miRNAs the most important miRNAs were Vun-miR395, Vun-miR396, Vun-miR390, Vun-miR393 and Vun-miR172 since they are connected 26, 16, 12, 12 and 9 times respectively with the VuRNAi genes (Fig 10 and supplementary file S5). During the development of plants, miRNAs are essential for controlling gene expression. For example, miR396 modulates innate immunity and grow defense against pathogen infection and it also regulates leaf development and phase change in *Arabidopsis* [109, 110]. In the literature it is found that Sulphur is an important secondary macronutrient that interacts with several stress regulating metabolites, hence improves plant growth, development and grain quality under various environmental stresses including drought and salinity. Sulfate allocation and accumulation are regulated by miR395 [111, 112]. When miR390 interacts with AGO7, it does so with extreme specificity, cleaving the targeted mRNA (106). Auxin is a phytohormone that regulates plant growth and development by binding on F-box (TAAR) receptors. Following pathogen infection, miR393 controls the expression of several sets of TAAR motifs, controls plant development and controls how the plant reacts to biotic and abiotic stressors [113, 114]. The miR172 family controls meristem size, trichome initiation, stem elongation, shoot branching, and floral fitness, with members exhibiting different expression patterns and functional specificity [115, 116]. Together, VuRNAi genes and miRNAs might function in the development of the plant’s defense mechanisms against biotic and abiotic stressors, as well as in its growth and development.

**Fig 10.**
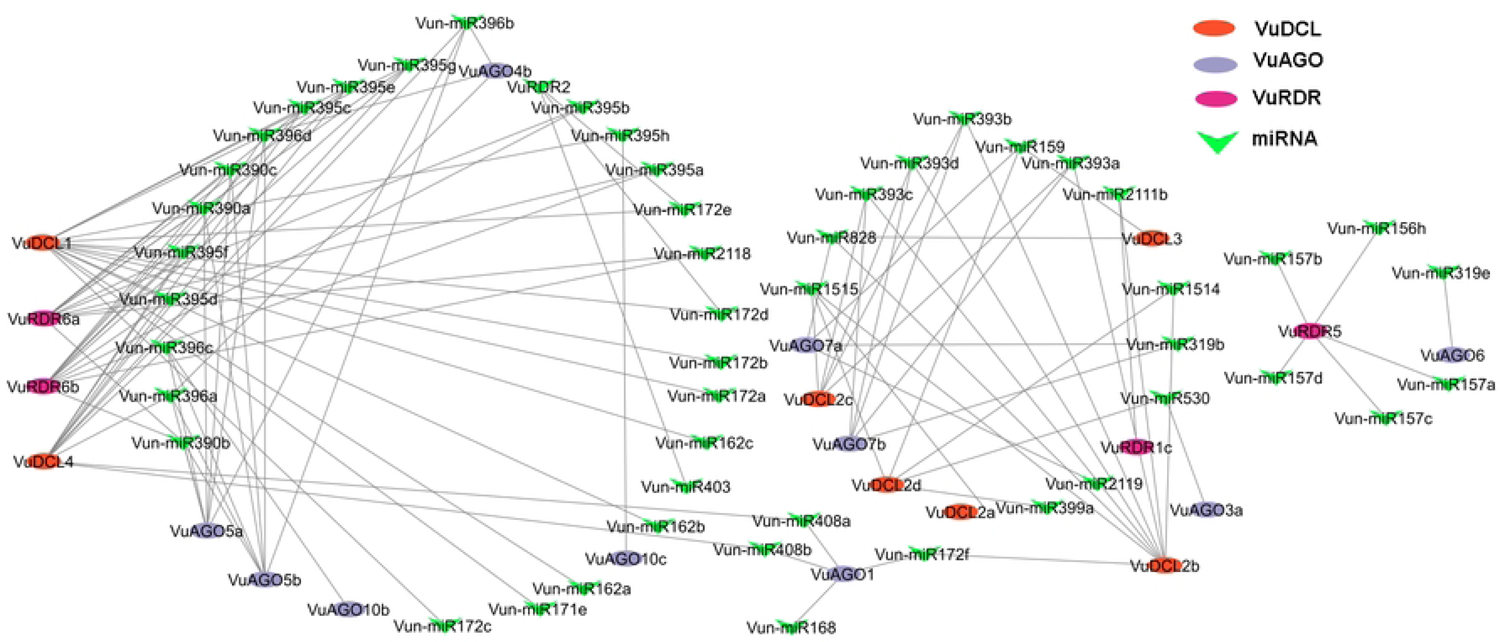
Regulatory network among the miRNAs and VuRNAi proteins.

#### 3.7.3 *Cis*-acting regulatory element analysis

The small motif (5–20 bp)-containing *cis*-acting elements (CAEs), which are often non-coding DNA parts positioned in the promoter region of target genes [117, 118]. The transcription of genes that bind to the target sites of CAEs is regulated by transcription factors (TFs) and transcriptional regulators (up- and down-regulators) [118]. We may now quickly access databases and estimate gene functions and their regulatory components inside their DNA sequence (usually the promoter and enhancer region), courtesy of the development of high-throughput genome sequencing technology. In this study, we used CAEs analysis from the online PLANTCARE database to identify the numerous significant motifs, their functional functions, and the diversity of the anticipated *VuDCL*, *VuAGO* and *VuRDR* genes (Fig 11; File S6). The CAEs were divided into the following five categories: 1) light responsive (LR), 2) stress responsive (SR), 3) hormone responsive (HR), 4) other activities (OT), and 5) unknown functions (UF). The first four categories were shown in Fig. 11, while the complete categories—including the unidentified functions—were provided in the supplemental File S6. The most significant physiological process in plants is photosynthesis, which takes place in the leaves’ tissue and is controlled by LR CAEs. The cowpea RNAi gene family’s upstream regulatory region has a higher density of LR elements among the CAEs (Fig 11). The majority of the RNAi-related proteins in cowpea had the LR elements ACE, AE-box, ATCT-motif, box-4, G-box, GATA-motif, GT1-motif, I-box, MRE, TCCC-motif, TCT-motif, and ATC-motif. We can infer from the previous work that the predicted LR motifs have a role in the light responsiveness of various plants [117,119–121]. As a result, the LR motif in cowpea has a considerable impact on the rate of photosynthesis in cowpea leaves and helps to improve grain quality and yield. Contrarily, hormones play a role in the growth and development of plants [122, 123]. The key cowpea HR motifs are ABRE (found in all cowpea RNAi genes), AuxRR-core, GARE-motif, P-box, TATC-box, TCA-element, and TGA-element HR CAEs shared by VuDCL, VuAGO, and *VuRDR* as phytohormone responsive elements [124–126] (Fig 11). Following this, it was discovered that the VuDCL, VuAGO, and VuRDR protein families of cowpea contain SR cis-acting elements that include TC-rich repeats involved in defense and stress responsiveness, MBS involved in drought inducibility and LTR motifs [127, 128]. There were also some other significantly important motifs like ARE, AT-rich element, CAAT-box, CCAAT-box, CGTCA-motif, GCN4-motif, circadian and CAT-box considered as other regulatory CAEs perform other functions [38, 43]. For example, the CAEs CAAT-box and TATA-box are the eukaryotic promoters [73] present in all RNAi proteins in cowpea. TGACG-motif presents in all RNAi family members related to salicylic acid responsive. CGTCA-motif related to Methyle Jasmonate (MeJA) regulating CAE present in cowpea. Along with reported CAEs some other unknown elements were predicted (File S6) in cowpea. Lastly, the CAEs that the cowpea projected RNAi gene family share can reveal crucial details about their functional capacity for plant growth, development, disease control and these CAEs could be exploited in building stress and disease resistance in plants, including cowpea.

**Fig 11.**
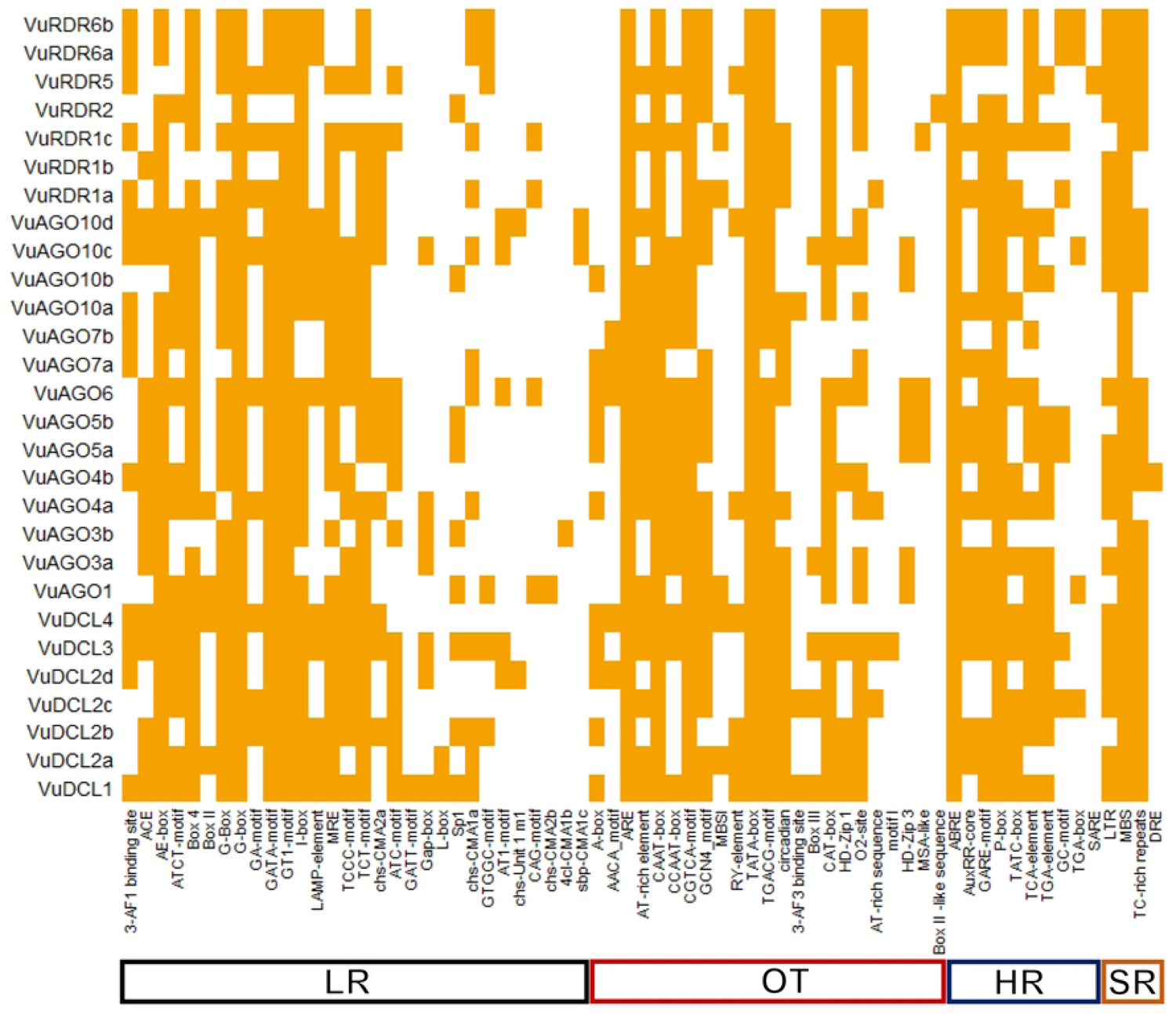
The expected *cis*-acting elements in the promoter region of the predicted VuRNAi genes, respectably. The deep color represents the presence of elements in the respective genes and white color represents absence of the elements in the respective genes.

## 4 Conclusion

Cowpea (*Vigna unguiculata* (L.) Walp) is a highly protein reached vegetable and food crops reduces malnutrition in impoverished countries as well as brings some health benefits like reducing the incidence of cardiovascular disease and some types of cancer. However, severe losses in cowpea production were observed around the globe due to different biotic and abiotic stresses like pathogens, droughts and salinity. Two 21–24 nucleotides long sRNAs: miRNA and siRNA regulate both TGS and PTGS ensuing plants growth, development, enhance natural immunity and grow resistance against biotic and abiotic stresses. The RNAi proteins encoded by RNAi genes play key role in the sRNAs biogenesis and functional processes. Nevertheless, so far, we know no studies have been done on the RNAi genes/proteins in cowpea. Therefore, in this study, we used a thorough computer-based *in silico* analysis to find and describe RNAi (AGO, DCL, and RDR) gene families across the whole cowpea genome while highlighting their related regulatory elements. We identified 28 VuRNAi genes including 7 *VuDCL*, 14 *VuAGO* and 7 *VuRDR* genes in cowpea. For identification of these VuRNAi genes in cowpea we conducted phylogenic, domain, motif, and gene structure analysis and compared them to AtRNAi genes to validate our prediction. To understand how VuRNAi genes contribute to plant growth, development and stress defense, we examined their chromosomal position, sub-cellular location, GO, TF, miRNA, and cis-regulatory elements. VuDCL1 may play a role in growth, stress, and flowering. We presume that VuDCL2 (a-d) contributes to the growth of the vegetative phase, disease resistance, and flowering. According to the results, *VuAGO7* and *VuAGO10* might be essential for plant growth. *VuRDR1* is increased antiviral defense, and transgenic silencing in cowpea. According to AtRDR6’s function, *VuRDR6* might create ta-siRNA precursor and degrade mRNA for antiviral defense. GO analysis suggests that VuRNAi genes participate in PTGS by cleaving mRNA in cowpea and stimulate the immune system. PAZ-Argonaute and SNF2 control *VuDCL* and *VuAGO*, both TFs involved in gene silencing. Additionally, miRNAs also help plants grow, develop, and deal with stress and they control how genes are expressed at both the transcriptional and posttranscriptional levels. Among the predicted miRNAs in cowpea miR395 controls sulfate accumulation and allocation in plants including cowpea, resultantly interacts with several stress regulating metabolites, hence improves plant growth, development and grain quality under various environmental stresses including drought and salinity. miR396 modifies immunity and defense against pathogen infection and regulates leaf development and phase change in Arabidopsis. The important predicted CAEs were LR, SR and HR *might be* involved in plant growth, development, improving production and grain quality and defense plants from biotic and abiotic stresses. Therefore, this study provides decisive information for further rigorous study regarding functional roles of VuRNAi gene and their regulatory elements for future abiotic and biotic stress resistance more robust cowpea.

### Supporting information

**S1 File:** Full-length protein sequences of DCL gene of *Vigna unguiculata* (L.) Walp. (TXT)

**S2 File:** Full-length protein sequences of AGO gene of *Vigna unguiculata* (L.) Walp. (TXT)

**S3 File:** Full-length protein sequences of RDR gene of *Vigna unguiculata* (L.) Walp. (TXT)

**S4 File:** Detail GO analysis results of the predicted VuRNAi genes. (XLSX)

**S5 File:** VuRNAi genes and associated *miRNA* analysis. (XLSX)

**S6 File:** VuRNAi genes and associated *cis*-acting regulatory elements analysis. (XLSX)

## Acknowledgments

The authors acknowledge ministry of science and technology, Bangladesh for the financial support, Bioinformatics Lab, Department of Statistics, University of Rajshahi, Bangladesh for the technical support and Department of Statistics, Bangabandhu Sheikh Mujibur Rahman Agricultural University, Bangladesh for the laboratory assistance.

## Author Contributions

**Conceptualization:** Mohammad Nazmol Hasan, Md Parvez Mosharaf, Darun Naim **Data curation:** Mohammad Nazmol Hasan, Keya Rani Das, Mst. Noorunnahar **Formal analysis:** Mohammad Nazmol Hasan **Supervision:** Md. Nurul Haque Mollah, Mohammad Nazmol Hasan **Visualization:** Mohammad Nazmol Hasan **Writing – original draft:** Mohammad Nazmol Hasan **Writing – review & editing:** Mohammad Nazmol Hasan, Md Parvez Mosharaf, Khandoker Saif Uddin, Keya Rani Das, Nasrin Sultana, Mst. Noorunnahar, Darun Naim and Md. Nurul Haque Mollah

## References

1. Burridge JD, Schneider HM, Huynh BL, Roberts PA, Bucksch A, Lynch JP. Genome-wide association mapping and agronomic impact of cowpea root architecture. Theor Appl Genet. 2017;130. doi:10.1007/s00122-016-2823-y

2. Huynh B, Close TJ, Roberts PA, Hu Z, Wanamaker S, Lucas MR, et al. Gene Pools and the Genetic Architecture of Domesticated Cowpea. Plant Genome. 2013;6. doi:10.3835/plantgenome2013.03.0005

3. Vasconcelos IM, Maia FMM, Farias DF, Campello CC, Carvalho AFU, de Azevedo Moreira R, et al. Protein fractions, amino acid composition and antinutritional constituents of high-yielding cowpea cultivars. J Food Compos Anal. 2010;23. doi:10.1016/j.jfca.2009.05.008

4. Jayathilake C, Visvanathan R, Deen A, Bangamuwage R, Jayawardana BC, Nammi S, et al. Cowpea: an overview on its nutritional facts and health benefits. Journal of the Science of Food and Agriculture. 2018. doi:10.1002/jsfa.9074

5. Adjei-Fremah S, Worku M, De Erive MO, He F, Wang T, Chen G. Effect of microfluidization on microstructure, protein profile and physicochemical properties of whole cowpea flours. Innov Food Sci Emerg Technol. 2019;57. doi:10.1016/j.ifset.2019.102207

6. Carneiro da Silva A, de Freitas Barbosa M, Bento da Silva P, Peres de Oliveira J, Loureiro da Silva T, Lopes Teixeira Junior D, et al. Health Benefits and Industrial Applications of Functional Cowpea Seed Proteins. Grain and Seed Proteins Functionality. 2021. doi:10.5772/intechopen.96257

7. Elharadallou SB, Khalid II, Gobouri AA, Abdel-Hafez SH. Amino Acid Composition of Cowpea (<i>Vigna ungiculata</i> L. Walp) Flour and Its Protein Isolates. Food Nutr Sci. 2015;06. doi:10.4236/fns.2015.69082

8. Carvalho M, Castro I, Moutinho-Pereira J, Correia C, Egea-Cortines M, Matos M, et al. Evaluating stress responses in cowpea under drought stress. J Plant Physiol. 2019;241. doi:10.1016/j.jplph.2019.153001

9. Loebenstein G, Thottappilly G. Virus and Virus-like Diseases of Major Crops in Developing Countries. Virus and Virus-like Diseases of Major Crops in Developing Countries. 2003. doi:10.1007/978-94-007-0791-7

10. Chen X. Small RNAs and their roles in plant development. Annual Review of Cell and Developmental Biology. 2009. doi:10.1146/annurev.cellbio.042308.113417

11. López-Gomollón S, Dalmay T. Recent patents in RNA silencing in plants: Constructs, methods and applications in plant biotechnology. Recent Patents DNA Gene Seq. 2010;4. doi:10.2174/187221510794751677

12. Sarkies P, Miska EA. Small RNAs break out: The molecular cell biology of mobile small RNAs. Nature Reviews Molecular Cell Biology. 2014. doi:10.1038/nrm3840

13. Voinnet O. Use, tolerance and avoidance of amplified RNA silencing by plants. Trends in Plant Science. 2008. doi:10.1016/j.tplants.2008.05.004

14. Balassa G, Balassa K, Janda T, Rudnóy S. Expression Pattern of RNA Interference Genes During Drought Stress and MDMV Infection in Maize. J Plant Growth Regul. 2022;41: 2048–2058. doi:10.1007/s00344-022-10651-z

15. Bharathi JK, Anandan R, Benjamin LK, Muneer S, Prakash MAS. Recent trends and advances of RNA interference (RNAi) to improve agricultural crops and enhance their resilience to biotic and abiotic stresses. Plant Physiol Biochem. 2023;194: 600–618. doi:10.1016/j.plaphy.2022.11.035

16. Lai EC. microRNAs: Runts of the Genome Assert Themselves. Current Biology. 2003. doi:10.1016/j.cub.2003.11.017

17. Finnegan EJ, Matzke MA. The small RNA world. J Cell Sci. 2003;116. doi:10.1242/jcs.00838

18. Shabalina SA, Koonin E V. Origins and evolution of eukaryotic RNA interference. Trends in Ecology and Evolution. 2008. doi:10.1016/j.tree.2008.06.005

19. Castel SE, Martienssen RA. RNA interference in the nucleus: Roles for small RNAs in transcription, epigenetics and beyond. Nature Reviews Genetics. 2013. doi:10.1038/nrg3355

20. Wilson RC, Doudna JA. Molecular mechanisms of RNA interference. Annu Rev Biophys. 2013;42. doi:10.1146/annurev-biophys-083012-130404

21. Valencia-Sanchez MA, Liu J, Hannon GJ, Parker R. Control of translation and mRNA degradation by miRNAs and siRNAs. Genes and Development. 2006. doi:10.1101/gad.1399806

22. Axtell MJ. Classification and comparison of small RNAs from plants. Annual Review of Plant Biology. 2013. doi:10.1146/annurev-arplant-050312-120043

23. Liu Q, Feng Y, Zhu Z. Dicer-like (DCL) proteins in plants. Functional and Integrative Genomics. 2009. doi:10.1007/s10142-009-0111-5

24. Carbonell A. Plant ARGONAUTES: Features, functions, and unknowns. Methods in Molecular Biology. 2017. doi:10.1007/978-1-4939-7165-7_1

25. Meister G, Tuschl T. Mechanisms of gene silencing by double-stranded RNA. Nature. 2004;431: 343–349. doi:10.1038/nature02873

26. Fagard M, Boutet S, Morel JB, Bellini C, Vaucheret H. AGO1, QDE-2, and RDE-1 are related proteins required for post-transcriptional gene silencing in plants, quelling in fungi, and RNA interference in animals. Proc Natl Acad Sci U S A. 2000;97. doi:10.1073/pnas.200217597

27. Zilberman D, Cao X, Jacobsen SE. ARGONAUTE4 control of locus-specific siRNA accumulation and DNA and histone methylation. Science (80-). 2003;299. doi:10.1126/science.1079695

28. Hunter C, Sun H, Poethig RS. The Arabidopsis Heterochronic Gene ZIPPY Is an ARGONAUTE Family Member. Curr Biol. 2003;13. doi:10.1016/j.cub.2003.09.004

29. Moussian B, Schoof H, Haecker A, Jürgens G, Laux T. Role of the ZWILLE gene in the regulation of central shoot meristem cell fate during Arabidopsis embryogenesis. EMBO J. 1998;17. doi:10.1093/emboj/17.6.1799

30. Hunter LJR, Brockington SF, Murphy AM, Pate AE, Gruden K, Macfarlane SA, et al. RNA-dependent RNA polymerase 1 in potato (Solanum tuberosum) and its relationship to other plant RNA-dependent RNA polymerases. Sci Rep. 2016;6. doi:10.1038/srep23082

31. Vazquez F. Arabidopsis endogenous small RNAs: highways and byways. Trends in Plant Science. 2006. doi:10.1016/j.tplants.2006.07.006

32. Kapoor M, Arora R, Lama T, Nijhawan A, Khurana JP, Tyagi AK, et al. Genome-wide identification, organization and phylogenetic analysis of Dicer-like, Argonaute and RNA-dependent RNA Polymerase gene families and their expression analysis during reproductive development and stress in rice. BMC Genomics. 2008;9. doi:10.1186/1471-2164-9-451

33. Qian Y, Cheng Y, Cheng X, Jiang H, Zhu S, Cheng B. Identification and characterization of Dicer-like, Argonaute and RNA-dependent RNA polymerase gene families in maize. Plant Cell Rep. 2011;30. doi:10.1007/s00299-011-1046-6

34. Bai M, Yang GS, Chen WT, Mao ZC, Kang HX, Chen GH, et al. Genome-wide identification of Dicer-like, Argonaute and RNA-dependent RNA polymerase gene families and their expression analyses in response to viral infection and abiotic stresses in Solanum lycopersicum. Gene. 2012;501. doi:10.1016/j.gene.2012.02.009

35. Yadav CB, Muthamilarasan M, Pandey G, Prasad M. Identification, characterization and expression profiling of dicer-like, argonaute and rna-dependent RNA polymerase gene families in foxtail millet. Plant Mol Biol Report. 2015;33. doi:10.1007/s11105-014-0736-y

36. Gan D, Zhan M, Yang F, Zhang Q, Hu K, Xu W, et al. Expression analysis of argonaute, Dicer-like, and RNA-dependent RNA polymerase genes in cucumber (Cucumis sativus L.) in response to abiotic stress. J Genet. 2017;96. doi:10.1007/s12041-017-0758-y

37. Qin L, Mo N, Muhammad T, Liang Y. Genome-wide analysis of DCL, AGO, and RDR gene families in pepper (Capsicum Annuum L.). Int J Mol Sci. 2018;19. doi:10.3390/ijms19041038

38. Mosharaf MP, Rahman H, Ahsan MA, Akond Z, Ahmed FF, Islam MM, et al. In silico identification and characterization of AGO, DCL and RDR gene families and their associated regulatory elements in sweet orange (Citrus sinensis L.). PLoS One. 2021;15. doi:10.1371/journal.pone.0228233

39. Goodstein DM, Shu S, Howson R, Neupane R, Hayes RD, Fazo J, et al. Phytozome: A comparative platform for green plant genomics. Nucleic Acids Res. 2012;40. doi:10.1093/nar/gkr944

40. Lonardi S, Muñoz-Amatriaín M, Liang Q, Shu S, Wanamaker SI, Lo S, et al. The genome of cowpea (Vigna unguiculata [L.] Walp.). Plant J. 2019;98: 767–782. doi:10.1111/tpj.14349

41. Altschul SF, Gish W, Miller W, Myers EW, Lipman DJ. Basic local alignment search tool. J Mol Biol. 1990;215. doi:10.1016/S0022-2836(05)80360-2

42. Lamesch P, Berardini TZ, Li D, Swarbreck D, Wilks C, Sasidharan R, et al. The Arabidopsis Information Resource (TAIR): Improved gene annotation and new tools. Nucleic Acids Res. 2012;40: 1202–1210. doi:10.1093/nar/gkr1090

43. Ahmed FF, Hossen MI, Sarkar MAR, Konak JN, Zohra FT, Shoyeb M, et al. Genome-wide identification of DCL, AGO and RDR gene families and their associated functional regulatory elements analyses in banana (Musa acuminata). PLoS One. 2021;16. doi:10.1371/journal.pone.0256873

44. Artimo P, Jonnalagedda M, Arnold K, Baratin D, Csardi G, De Castro E, et al. ExPASy: SIB bioinformatics resource portal. Nucleic Acids Res. 2012;40. doi:10.1093/nar/gks400

45. Kumar S, Stecher G, Tamura K. MEGA7: Molecular Evolutionary Genetics Analysis Version 7.0 for Bigger Datasets. Mol Biol Evol. 2016;33. doi:10.1093/MOLBEV/MSW054

46. Mistry J, Chuguransky S, Williams L, Qureshi M, Salazar GA, Sonnhammer ELL, et al. Pfam: The protein families database in 2021. Nucleic Acids Res. 2021;49. doi:10.1093/nar/gkaa913

47. Bailey TL, Johnson J, Grant CE, Noble WS. The MEME Suite. Nucleic Acids Res. 2015;43. doi:10.1093/nar/gkv416

48. Hu B, Jin J, Guo AY, Zhang H, Luo J, Gao G. GSDS 2.0: An upgraded gene feature visualization server. Bioinformatics. 2015;31. doi:10.1093/bioinformatics/btu817

49. Liu L, Zhang Z, Mei Q, Chen M. PSI: A Comprehensive and Integrative Approach for Accurate Plant Subcellular Localization Prediction. PLoS One. 2013;8. doi:10.1371/journal.pone.0075826

50. Tian T, Liu Y, Yan H, You Q, Yi X, Du Z, et al. AgriGO v2.0: A GO analysis toolkit for the agricultural community, 2017 update. Nucleic Acids Res. 2017;45. doi:10.1093/nar/gkx382

51. Lescot M, Déhais P, Thijs G, Marchal K, Moreau Y, Van De Peer Y, et al. PlantCARE, a database of plant cis-acting regulatory elements and a portal to tools for in silico analysis of promoter sequences. Nucleic Acids Res. 2002;30. doi:10.1093/nar/30.1.325

52. Großhans H, Filipowicz W. The expanding world of small RNAs. Nature. 2008;451. doi:10.1038/451414a

53. Millar AA, Waterhouse PM. Plant and animal microRNAs: Similarities and differences. Functional and Integrative Genomics. 2005. doi:10.1007/s10142-005-0145-2

54. Bologna NG, Voinnet O. The diversity, biogenesis, and activities of endogenous silencing small RNAs in Arabidopsis. Annual Review of Plant Biology. 2014. doi:10.1146/annurev-arplant-050213-035728

55. Fei Q, Xia R, Meyers BC. Phased, secondary, small interfering RNAs in posttranscriptional regulatory networks. Plant Cell. 2013. doi:10.1105/tpc.113.114652

56. Holoch D, Moazed D. RNA-mediated epigenetic regulation of gene expression. Nature Reviews Genetics. 2015. doi:10.1038/nrg3863

57. Schmitz RJ, Tamada Y, Doyle MR, Zhang X, Amasino RM. Histone H2B deubiquitination is required for transcriptional activation of Flowering Locus C and for proper control of flowering in Arabidopsis. Plant Physiol. 2009;149. doi:10.1104/pp.108.131508

58. Sahu PP, Sharma N, Puranik S, Prasad M. Post-transcriptional and Epigenetic Arms of RNA Silencing: A Defense Machinery of Naturally Tolerant Tomato Plant Against Tomato Leaf Curl New Delhi Virus. Plant Mol Biol Report. 2014;32. doi:10.1007/s11105-014-0708-2

59. Henderson IR, Zhang X, Lu C, Johnson L, Meyers BC, Green PJ, et al. Dissecting Arabidopsis thaliana DICER function in small RNA processing, gene silencing and DNA methylation patterning. Nat Genet. 2006;38. doi:10.1038/ng1804

60. Carmell MA, Xuan Z, Zhang MQ, Hannon GJ. The Argonaute family: Tentacles that reach into RNAi, developmental control, stem cell maintenance, and tumorigenesis. Genes and Development. 2002. doi:10.1101/gad.1026102

61. Vaucheret H, Vazquez F, Crété P, Bartel DP. The action of ARGONAUTE1 in the miRNA pathway and its regulation by the miRNA pathway are crucial for plant development. Genes Dev. 2004;18: 1187–1197. doi:10.1101/gad.1201404

62. Zilberman D, Cao X, Johansen LK, Xie Z, Carrington JC, Jacobsen SE. Role of Arabidopsis ARGONAUTE4 in RNA-directed DNA methylation triggered by inverted repeats. Curr Biol. 2004;14. doi:10.1016/j.cub.2004.06.055

63. Lynn K, Fernandez A, Aida M, Sedbrook J, Tasaka M, Masson P, et al. The PINHEAD/ZWILLE gene acts pleiotropically in Arabidopsis development and has overlapping functions with the ARGONAUTE1 gene. Development. 1999;126. doi:10.1242/dev.126.3.469

64. Schiebel W, Pelissier T, Riedel L, Thalmeir S, Schiebel R, Kempe D, et al. Isolation of an RNA-Directed RNA Polymerase-Specific cDNA Clone from Tomato. Plant Cell. 1998;10. doi:10.2307/3870786

65. Sijen T, Fleenor J, Simmer F, Thijssen KL, Parrish S, Timmons L, et al. On the role of RNA amplification in dsRNA-triggered gene silencing. Cell. 2001;107. doi:10.1016/S0092-8674(01)00576-1

66. Jovel J, Walker M, Sanfaçon H. Recovery of Nicotiana benthamiana Plants from a Necrotic Response Induced by a Nepovirus Is Associated with RNA Silencing but Not with Reduced Virus Titer. J Virol. 2007;81. doi:10.1128/jvi.01192-07

67. Yu D, Fan B, MacFarlane SA, Chen Z. Analysis of the involvement of an inducible Arabidopsis RNA-dependent RNA polymerase in antiviral defense. Mol Plant-Microbe Interact. 2003;16. doi:10.1094/MPMI.2003.16.3.206

68. Chan SWL, Zilberman D, Xie Z, Johansen LK, Carrington JC, Jacobsen SE. RNA Silencing Genes Control de Novo DNA Methylation. Science (80-). 2004;303. doi:10.1126/science.1095989

69. Xie Z, Fan B, Chen C, Chen Z. An important role of an inducible RNA-dependent RNA polymerase in plant antiviral defense. Proc Natl Acad Sci U S A. 2001;98. doi:10.1073/pnas.111440998

70. Xie Z, Johansen LK, Gustafson AM, Kasschau KD, Lellis AD, Zilberman D, et al. Genetic and functional diversification of small RNA pathways in plants. PLoS Biol. 2004;2. doi:10.1371/journal.pbio.0020104

71. Matzke M, Kanno T, Daxinger L, Huettel B, Matzke AJ. RNA-mediated chromatin-based silencing in plants. Current Opinion in Cell Biology. 2009. doi:10.1016/j.ceb.2009.01.025

72. Yoshikawa M, Peragine A, Mee YP, Poethig RS. A pathway for the biogenesis of trans-acting siRNAs in Arabidopsis. Genes Dev. 2005;19. doi:10.1101/gad.1352605

73. Cui DL, Meng JY, Ren XY, Yue JJ, Fu HY, Huang MT, et al. Genome-wide identification and characterization of DCL, AGO and RDR gene families in Saccharum spontaneum. Sci Rep. 2020;10. doi:10.1038/s41598-020-70061-7

74. MacRae IJ, Doudna JA. Ribonuclease revisited: structural insights into ribonuclease III family enzymes. Current Opinion in Structural Biology. 2007. doi:10.1016/j.sbi.2006.12.002

75. Katsarou K, Mavrothalassiti E, Dermauw W, Van Leeuwen T, Kalantidis K. Combined Activity of DCL2 and DCL3 Is Crucial in the Defense against Potato Spindle Tuber Viroid. PLoS Pathog. 2016;12. doi:10.1371/journal.ppat.1005936

76. Moazed D. Small RNAs in transcriptional gene silencing and genome defence. Nature. 2009. doi:10.1038/nature07756

77. Hutvagner G, Simard MJ. Argonaute proteins: Key players in RNA silencing. Nature Reviews Molecular Cell Biology. 2008. doi:10.1038/nrm2321

78. Simon B, Kirkpatrick JP, Eckhardt S, Reuter M, Rocha EA, Andrade-Navarro MA, et al. Recognition of 2′-o-methylated 3′-end of piRNA by the PAZ domain of a Piwi protein. Structure. 2011;19. doi:10.1016/j.str.2010.11.015

79. Arribas-Hernández L, Marchais A, Poulsen C, Haase B, Hauptmann J, Benes V, et al. The slicer activity of ARGONAUTE1 is required specifically for the phasing, not production, of trans-acting short interfering RNAs in arabidopsis. Plant Cell. 2016;28. doi:10.1105/tpc.16.00121

80. Fang X, Qi Y. Rnai in plants: An argonaute-centered view. Plant Cell. 2015. doi:10.1105/tpc.15.00920

81. Höck J, Meister G. The Argonaute protein family. Genome Biology. 2008. doi:10.1186/gb-2008-9-2-210

82. Rivas F V., Tolia NH, Song JJ, Aragon JP, Liu J, Hannon GJ, et al. Purified Argonaute2 and an siRNA form recombinant human RISC. Nat Struct Mol Biol. 2005;12. doi:10.1038/nsmb918

83. Chung BYW, Valli A, Deery MJ, Navarro FJ, Brown K, Hnatova S, et al. Distinct roles of Argonaute in the green alga Chlamydomonas reveal evolutionary conserved mode of miRNA-mediated gene expression. Sci Rep. 2019;9. doi:10.1038/s41598-019-47415-x

84. Wassenegger M, Krczal G. Nomenclature and functions of RNA-directed RNA polymerases. Trends in Plant Science. 2006. doi:10.1016/j.tplants.2006.01.003

85. Venkataraman S, Prasad BVLS, Selvarajan R. RNA dependent RNA polymerases: Insights from structure, function and evolution. Viruses. 2018. doi:10.3390/v10020076

86. Marker S, Le Mouël A, Meyer E, Simon M. Distinct RNA-dependent RNA polymerases are required for RNAi triggered by double-stranded RNA versus truncated transgenes in Paramecium tetraurelia. Nucleic Acids Res. 2010;38. doi:10.1093/nar/gkq131

87. Chen C, Chen H, Zhang Y, Thomas HR, Frank MH, He Y, et al. TBtools: An Integrative Toolkit Developed for Interactive Analyses of Big Biological Data. Mol Plant. 2020;13. doi:10.1016/j.molp.2020.06.009

88. Arumuganathan K, Earle ED. Nuclear DNA content of some important plant species. Plant Mol Biol Report. 1991;9. doi:10.1007/BF02672069

89. Hoffer P, Ivashuta S, Pontes O, Vitins A, Pikaard C, Mroczka A, et al. Posttranscriptional gene silencing in nuclei. Proc Natl Acad Sci U S A. 2011;108. doi:10.1073/pnas.1009805108

90. Agrawal N, Dasaradhi PVN, Mohmmed A, Malhotra P, Bhatnagar RK, Mukherjee SK. RNA Interference: Biology, Mechanism, and Applications. Microbiol Mol Biol Rev. 2003;67. doi:10.1128/mmbr.67.4.657-685.2003

91. Chery J. RNA therapeutics: RNAi and antisense mechanisms and clinical applications. Postdoc J. 2016;4. doi:10.14304/surya.jpr.v4n7.5

92. Ehrlich JS, Hansen MDH, Nelson WJ. Spatio-temporal regulation of Rac1 localization and lamellipodia dynamics during epithelial cell-cell adhesion. Dev Cell. 2002;3. doi:10.1016/S1534-5807(02)00216-2

93. Glory E, Murphy RF. Automated Subcellular Location Determination and High-Throughput Microscopy. Developmental Cell. 2007. doi:10.1016/j.devcel.2006.12.007

94. Lingel A, Izaurralde E. RNAi: Finding the elusive endonuclease. RNA. 2004. doi:10.1261/rna.7175704

95. Fustin JM, Doi M, Yamaguchi Y, Hida H, Nishimura S, Yoshida M, et al. XRNA-methylation-dependent RNA processing controls the speed of the circadian clock. Cell. 2013;155: 793. doi:10.1016/j.cell.2013.10.026

96. Cui J, Liu RD, Wang L, Zhang X, Jiang P, Liu MY, et al. Proteomic analysis of surface proteins of Trichinella spiralis muscle larvae by two-dimensional gel electrophoresis and mass spectrometry. Parasites and Vectors. 2013;6. doi:10.1186/1756-3305-6-355

97. Fire A, Xu S, Montgomery MK, Kostas SA, Driver SE, Mello CC. Potent and specific genetic interference by double-stranded RNA in caenorhabditis elegans. Nature. 1998;391. doi:10.1038/35888

98. Shu Y, Liu Y, Zhang J, Song L, Guo C. Genome-wide analysis of the AP2/ERF superfamily genes and their responses to abiotic stress in Medicago truncatula. Front Plant Sci. 2016;6. doi:10.3389/fpls.2015.01247

99. Khan SA, Li MZ, Wang SM, Yin HJ. Revisiting the role of plant transcription factors in the battle against abiotic stress. International Journal of Molecular Sciences. 2018. doi:10.3390/ijms19061634

100. Sasaki K. Utilization of transcription factors for controlling floral morphogenesis in horticultural plants. Breeding Science. 2018. doi:10.1270/jsbbs.17114

101. Latchman DS. Transcription factors: An overview. Int J Biochem Cell Biol. 1997;29. doi:10.1016/S1357-2725(97)00085-X

102. Baumberger N, Baulcombe DC. Arabidopsis ARGONAUTE1 is an RNA Slicer that selectively recruits microRNAs and short interfering RNAs. Proc Natl Acad Sci U S A. 2005;102. doi:10.1073/pnas.0505461102

103. Carbonell A, Fahlgren N, Garcia-Ruiz H, Gilbert KB, Montgomery TA, Nguyen T, et al. Functional analysis of three Arabidopsis argonautes using slicer-defective mutants. Plant Cell. 2012;24. doi:10.1105/tpc.112.099945

104. Neves-Costa A, Will WR, Vetter AT, Miller JR, Varga-Weisz P. The SNF2-family member FUN30 promotes gene silencing in heterochromatic loci. PLoS One. 2009;4. doi:10.1371/journal.pone.0008111

105. Knizewski L, Ginalski K, Jerzmanowski A. Snf2 proteins in plants: gene silencing and beyond. Trends in Plant Science. 2008. doi:10.1016/j.tplants.2008.08.004

106. Katsarou K, Mitta E, Bardani E, Oulas A, Dadami E, Kalantidis K. DCL-suppressed Nicotiana benthamiana plants: valuable tools in research and biotechnology. Mol Plant Pathol. 2019;20. doi:10.1111/mpp.12761

107. Bartel DP. MicroRNAs: Genomics, Biogenesis, Mechanism, and Function. Cell. 2004. doi:10.1016/S0092-8674(04)00045-5

108. Faraoni I, Antonetti FR, Cardone J, Bonmassar E. miR-155 gene: A typical multifunctional microRNA. Biochimica et Biophysica Acta - Molecular Basis of Disease. 2009. doi:10.1016/j.bbadis.2009.02.013

109. Hou N, Cao Y, Li F, Yuan W, Bian H, Wang J, et al. Epigenetic regulation of miR396 expression by SWR1-C and the effect of miR396 on leaf growth and developmental phase transition in Arabidopsis. J Exp Bot. 2019;70: 5217–5229. doi:10.1093/jxb/erz285

110. Soto-Suárez M, Baldrich P, Weigel D, Rubio-Somoza I, San Segundo B. The Arabidopsis miR396 mediates pathogen-associated molecular pattern-triggered immune responses against fungal pathogens. Sci Rep. 2017;7: 1–14. doi:10.1038/srep44898

111. Liang G, Yang F, Yu D. MicroRNA395 mediates regulation of sulfate accumulation and allocation in Arabidopsis thaliana. Plant J. 2010;62. doi:10.1111/j.1365-313X.2010.04216.x

112. Kawashima CG, Yoshimoto N, Maruyama-Nakashita A, Tsuchiya YN, Saito K, Takahashi H, et al. Sulphur starvation induces the expression of microRNA-395 and one of its target genes but in different cell types. Plant J. 2009;57. doi:10.1111/j.1365-313X.2008.03690.x

113. Jiang J, Zhu H, Li N, Batley J, Wang Y. The miR393-Target Module Regulates Plant Development and Responses to Biotic and Abiotic Stresses. Int J Mol Sci. 2022;23. doi:10.3390/ijms23169477

114. Si-Ammour A, Windels D, Arn-Bouldoires E, Kutter C, Ailhas J, Meins F, et al. miR393 and secondary siRNAs regulate expression of the TIR1/AFB2 auxin receptor clade and auxin-related development of Arabidopsis leaves. Plant Physiol. 2011;157: 683–691. doi:10.1104/pp.111.180083

115. Lian H, Wang L, Ma N, Zhou CM, Han L, Zhang TQ, et al. Redundant and specific roles of individual MIR172 genes in plant development. PLoS Biol. 2021;19: 1–25. doi:10.1371/journal.pbio.3001044

116. Zhu QH, Helliwell CA. Regulation of flowering time and floral patterning by miR172. J Exp Bot. 2011;62: 487–495. doi:10.1093/jxb/erq295

117. Kaur A, Pati PK, Pati AM, Nagpal AK. In-silico analysis of cis-acting regulatory elements of pathogenesis-related proteins of Arabidopsis thaliana and Oryza sativa. PLoS One. 2017;12. doi:10.1371/journal.pone.0184523

118. Wittkopp PJ, Kalay G. Cis-regulatory elements: Molecular mechanisms and evolutionary processes underlying divergence. Nature Reviews Genetics. 2012. doi:10.1038/nrg3095

119. Ishige F, Takaichi M, Foster R, Chua NH, Oeda K. A G-box motif (GCCACGTGCC) tetramer confers high-level constitutive expression in dicot and monocot plants. Plant J. 1999;18. doi:10.1046/j.1365-313X.1999.00456.x

120. Le Gourrierec J, Li YF, Zhou DX. Transcriptional activation by Arabidopsis GT-1 may be through interaction with TFIIA-TBP-TATA complex. Plant J. 1999;18. doi:10.1046/j.1365-313X.1999.00482.x

121. Menkens AE, Schindler U, Cashmore AR. The G-box: a ubiquitous regulatory DNA element in plants bound by the GBF family of bZIP proteins. Trends in Biochemical Sciences. 1995. doi:10.1016/S0968-0004(00)89118-5

122. Gray WM. Hormonal regulation of plant growth and development. PLoS Biology. 2004. doi:10.1371/journal.pbio.0020311

123. Ogawa M, Hanada A, Yamauchi Y, Kuwahara A, Kamiya Y, Yamaguchi S. Gibberellin biosynthesis and response during Arabidopsis seed germination. Plant Cell. 2003;15. doi:10.1105/tpc.011650

124. Maruyama K, Todaka D, Mizoi J, Yoshida T, Kidokoro S, Matsukura S, et al. Identification of cis-acting promoter elements in cold-and dehydration-induced transcriptional pathways in arabidopsis, rice, and soybean. DNA Res. 2012;19. doi:10.1093/dnares/dsr040

125. Ezcurra I, Wycliffe P, Nehlin L, Ellerström M, Rask L. Transactivation of the Brassica napus napin promoter by ABI3 requires interaction of the conserved B2 and B3 domains of ABI3 with different cis-elements: B2 mediates activation through an ABRE, whereas B3 interacts with an RY/G-box. Plant J. 2000;24. doi:10.1046/j.1365-313X.2000.00857.x

126. Zhou Y, Hu L, Wu H, Jiang L, Liu S. Genome-Wide Identification and Transcriptional Expression Analysis of Cucumber Superoxide Dismutase (SOD) Family in Response to Various Abiotic Stresses. Int J Genomics. 2017;2017. doi:10.1155/2017/7243973

127. Arias JA, Dixon RA, Lamb CJ. Dissection of the functional architecture of a plant defense gene promoter using a homologous in vitro transcription initiation system. Plant Cell. 1993;5. doi:10.1105/tpc.5.4.485

128. Chon W, Provart NJ, Glazebrook J, Katagiri F, Chang HS, Eulgem T, et al. Expression profile matrix of Arabidopsis transcription factor genes suggests their putative functions in response to environmental stresses. Plant Cell. 2002;14. doi:10.1105/tpc.010410

